# Comparing quality of reporting between preprints and peer-reviewed articles in the biomedical literature

**DOI:** 10.1101/581892

**Authors:** Clarissa F. D. Carneiro, Victor G. S. Queiroz, Thiago C. Moulin, Carlos A. M. Carvalho, Clarissa B. Haas, Danielle Rayêe, David E. Henshall, Evandro A. De-Souza, Felippe E. Amorim, Flávia Z. Boos, Gerson D. Guercio, Igor R. Costa, Karina L. Hajdu, Lieve van Egmond, Martin Modrák, Pedro B. Tan, Richard J. Abdill, Steven J. Burgess, Sylvia F. S. Guerra, Vanessa T. Bortoluzzi, Olavo B. Amaral

## Abstract

**Background:** Preprint usage is growing rapidly in the life sciences; however, questions remain on the relative quality of preprints when compared to published articles. An objective dimension of quality that is readily measurable is completeness of reporting, as transparency can improve the reader’s ability to independently interpret data and reproduce findings.

**Methods:** In this observational study, we initially compared independent samples of articles published in bioRxiv and in PubMed-indexed journals in 2016 using a quality of reporting questionnaire. After that, we performed paired comparisons between preprints from bioRxiv to their own peer-reviewed versions in journals.

**Results:** Peer-reviewed articles had, on average, higher quality of reporting than preprints, although the difference was small, with absolute differences of 5.0% [95% CI 1.4, 8.6] and 4.7% [95% CI 2.4, 7.0] of reported items in the independent samples and paired sample comparison, respectively. There were larger differences favoring peer-reviewed articles in subjective ratings of how clearly titles and abstracts presented the main findings and how easy it was to locate relevant reporting information. Changes in reporting from preprints to peer-reviewed versions did not correlate with the impact factor of the publication venue or with the time lag from bioRxiv to journal publication.

**Conclusions:** Our results suggest that, on average, publication in a peer-reviewed journal is associated with improvement in quality of reporting. They also show that quality of reporting in preprints in the life sciences is within a similar range as that of peer-reviewed articles, albeit slightly lower on average, supporting the idea that preprints should be considered valid scientific contributions.

## Introduction

Editorial peer review refers to the process whereby researchers from relevant fields review scientific articles with the purpose of evaluating their quality and/or adequacy to a publication venue. The debate on the origin of this practice revolves around how broadly it is defined; however, articles have been evaluated by various forms of peer review since the creation of scientific journals (for a historical review, see Csiszar, 2016).

Despite the ubiquity of editorial peer review, we have little empirical evidence supporting its effectiveness to ensure article quality (Jefferson et al., 2007). Evaluations limited to individual journals (Goodman et al., 1994; Pierie et al., 1996) have shown that peer review slightly improves reporting of various items, with the greatest improvements observed in the discussion and conclusion sections. Nevertheless, evaluations of its effect on research quality have not been performed in more representative samples of the literature. Moreover, positive effects of peer review in individual journals do not necessarily imply that it will work as an effective filter on a systemic level (Ioannidis et al., 2010).

Additionally, traditional peer review has various drawbacks (Walker and da Silva, 2015), including reviewer bias (Mahoney, 1977; Murray et al., 2018), lack of agreement among reviewers (Rothwell and Martyn, 2000; Pier et al., 2018) and vulnerability to various forms of system gaming such as ‘lottery behavior’ by authors (Ioannidis et al., 2010), predatory journals (Bohannon, 2013) and self-peer-review scams (Ferguson et al., 2014). Its most often quoted limitation, however, is the time lag for publication of articles (Vale, 2015; Berg et al., 2016; Cobb, 2017) and the resulting delay in the dissemination of scientific findings. Due to its gatekeeping function, editorial peer review has also become associated with other problems of scientific publication, such as paywalls and high prices imposed by commercial publishers. In view of these problems, various initiatives have tried to reform or bypass peer review in order to provide faster and wider access to scientific knowledge.

Preprints are complete manuscripts submitted to publicly accessible repositories, which may or may not later be submitted to a formal scientific journal. Preprint usage is common in communities such as physics and mathematics, particularly due to the popularity of arXiv, a seminal preprint server established in 1991 (Ginsparg, 2011). Spurred by the recent creation of new repositories such as bioRxiv and PeerJ, as well as by scientist-driven initiatives to support their use (Berg et al., 2016), biomedical scientists have recently become more adept at the practice (Cobb, 2017). Nevertheless, reward systems still largely rely on formal journal publication, leading to a dissociation between the dissemination of scientific findings through preprints from the certification provided by peer review (Cobb, 2017).

Predictably, the main concerns about this model of scientific communication revolve around the quality of non-peer-reviewed studies (Vale, 2015; Berg et al., 2016; Calne, 2016). At the same time, however, preprints offer a unique opportunity to study the effects of peer review, by allowing comparisons between non-reviewed manuscripts with their final published versions. Studies of samples from arXiv and bioRxiv using automated text measures have shown that changes from pre- to post-peer-review versions are usually minor (Klein et al., 2018). Nevertheless, to our knowledge, no attempt has been made to evaluate changes in study quality.

Scientific quality has many dimensions, such as rigor in methodological design, novelty and impact of findings, and transparency of reporting. Evaluating the appropriateness of methodology or the significance of results on a wide scale is challenging, due to the inherent subjectivity of these judgments and the need for area-specific expertise. Transparency and quality of reporting, however, can be assessed more objectively, with reporting guidelines and checklists available in many fields of science to guide authors on the minimum information that a manuscript should include (Simera et al., 2009). Quality of reporting is used to evaluate study quality in meta-analyses (Ryan et al., 2013), as well as the effect of interventions focused on improving transparency (Han et al., 2017; Hair, Macleod and Sena, 2019; The NPQIP Collaborative group, 2019). Moreover, it may be the aspect of manuscript quality that is most readily amenable to improvement by peer review, as reporting issues should be relatively simple to detect and fix.

In this study, we aim to compare quality of reporting between preprints and peer-reviewed articles in the life sciences. For this, we compiled a simplified list of essential items that should be reported in different types of biomedical articles, based on existing checklists (Moher, Schulz and Altman, 2001; von Elm et al., 2007; Kilkenny et al., 2010; Bossuyt et al., 2015; Hair, Macleod and Sena, 2019; The NPQIP Collaborative group, 2019). We first selected independent random samples of preprints from bioRxiv and peer-reviewed articles from PubMed, in order to compare quality of reporting between them. We then performed a paired comparison of a sample of preprints from bioRxiv to their own peer-reviewed versions in order to more directly assess the effects of peer review.

## Materials and Methods

Data collection and analysis protocols were preregistered for the comparison between bioRxiv and PubMed articles (hereby referred to as “independent samples comparison”) at https://osf.io/rxqn4. These were later updated at https://osf.io/g3ehr/ for the comparison between preprints and their published versions (hereby referred to as “paired sample comparison”). Analyses that were not included in the original plan will be referred to as exploratory throughout the text.

### Study selection

#### Independent samples comparison (bioRxiv vs. PubMed)

We obtained a list of all articles published in PubMed and bioRxiv between January 1st and December 31st, 2016. This date range had to comprise the first version of a preprint or the online publication date for peer-reviewed articles. Although we cannot be sure that the first preprint version had not undergone peer review before its publication, the most common practice seems to be to post a preprint before or at the moment of submission to a peer-reviewed journal (Sever et al., 2019). Articles were randomly selected using the *sample* function in R and were double-screened by the coordinating team (C.F.D.C, V.G.S.Q., T.C.M. or O.B.A.) for the following inclusion criteria: articles should i) be written in English, ii) contain at least one original result, iii) include a statistical comparison between different experimental or observational groups and iv) have groups composed of human or non-human animals, cells, microorganisms or biological samples derived from them. We selected the first result presented in each article that filled these criteria, consisting of a single figure/subpanel or table, which was then used for analysis. Disagreements on inclusion were discussed by the coordinating team until consensus was reached.

Articles were categorized according to the biological model (in vitro/cell lines, invertebrates, vertebrates and humans), and the number of articles per category was matched across groups. Thus, each selected study was included in the independent samples comparison according to the availability of selected studies in the other group until our planned sample size was reached.

#### Paired sample comparison (preprints vs. peer-reviewed versions)

Preprints selected by the process described above were later evaluated for inclusion in the paired sample if (i) their bioRxiv page listed a peer-reviewed publication, (ii) the date of publication was no later than December 31st, 2018 and (iii) the same figure/subpanel/table selected previously was present on the main text of the peer-reviewed publication.

### Data collection

#### Quality of reporting evaluation

Evaluation of each study was performed through an online questionnaire implemented on Google Forms. Questions were based on existing reporting guidelines (Moher et al., 2001; von Elm et al., 2007; Kilkenny et al., 2010; Bossuyt et al., 2015), journal checklists (Nature, 2013) and previous studies on quality of reporting (Hair, Macleod and Sena, 2019; The NPQIP Collaborative group, 2019), and are presented along with their response options on **Table S1**. They were based on direct, objective criteria, in an attempt to avoid the need for subjective evaluation. Analyzed reporting items included measures to reduce risk of bias (e.g. blinding, conflict of interest reporting), details on reagents (e.g. antibody validation, reagent source), data presentation (e.g. summary and variation measures, identifiable groups, definition of symbols used), data analysis (e.g. statistical tests used, exact p values) and details on the biological model (e.g. culture conditions, animal species and strain, human subject recruitment and eligibility, ethical requirements). As not all of these apply to every article, some questions were category-specific, while others could be answered as ‘not applicable’. A detailed Instructions Manual for answering the questions (available as **Supplementary Text 1**) was distributed to evaluators to standardize interpretation. Importantly, most questions concerned only the result selected for analysis (i.e. the first table, figure or subpanel fulfilling our inclusion criteria) and not the whole set of results.

Two additional questions regarding evaluators’ subjective assessments were included in the questionnaire, to be answered on a five-point scale. The first asked whether the title and abstract provided a clear idea of the article’s main findings, ranging from “Not clear at all” to “Perfectly clear”. The second one asked whether the information required in the questionnaire was easy to find and extract from the article, ranging from “Very hard” to “Very easy”.

Evaluators were biomedical researchers recruited locally at Brazilian universities and online through the ASAPbio blog (Amaral, 2018) and social media. To be included as evaluators, candidates had to reach an agreement of at least 75% in a test set of 4 articles. This comparison was based on the consensus answers of 3 members of the coordinating team (C.F.D.C, T.C.M. and O.B.A.) for 2 sets of 4 articles, reached after extensive discussion over possible disagreements. A candidate who failed to reach the required level of agreement on the first set could try again on the second set after reviewing his own answers along with the consensus in the first test. After achieving the agreement threshold, evaluators had access to the consensus answers as well as their own on the evaluated set(s).

As the paired sample comparison was started almost a year after the independent samples one, we sought to determine whether the initial analysis of preprints could be reused for the paired sample. For this, we performed correlations between time and score for each evaluator in the first stage and compared the mean r value to zero. Additionally, we performed equivalence tests between the score obtained in the first stage to the score from an independent reanalysis by a single evaluator in the second stage for a sample of 35 preprints. Though there was no clear evidence that individual evaluators changed their scoring over time, the equivalence test (with an estimated power of 90% to detect equivalence at ± 5% with α=0.05) failed to provide statistical evidence for equivalence at the ± 5% bound (see https://osf.io/g3ehr/ and https://osf.io/h7s3g/ for details). Therefore, all preprints included in the paired sample comparison were reanalyzed to avoid any time-related bias in the comparison between preprints and their published versions.

Each article was assessed independently by three evaluators, and the most prevalent answer among them for each question was considered final (except for subjective assessments, where the final score was the mean of the three evaluations). If all three evaluators reached different answers (a possibility arising when more than two response options were available), the question was discussed by the coordinating team until consensus was reached.

PDF files were redacted so that evaluators were blinded to the journal, list of authors, their affiliation and funders. However, some of this information could still be inferred from the formatting of the PDF file or from methodological details (such as the ethics committee or place of sample collection). As we considered typesetting to be a direct consequence of the editorial process, we chose to maintain the original formatting of articles, which meant that most journal articles were recognizable as such. Consequently, evaluators were not blinded to the group of origin of articles.

#### Reporting scores

The overall reporting scores were defined as the percentage of items reported for each article, using the total number of applicable questions – defined both by the biological model category and by the number of questions rated by the evaluators as not applicable – as the denominator. General reporting scores considered only the questions in the first five sections of the questionnaire, while specific scores considered the section for the corresponding biological model of the result under analysis. For some questions, a partial score was assigned for partial reporting, as described in **Table S1**.

#### Evaluator agreement

Agreement between individual pairs of evaluators was calculated as the mean percentage of identical responses between them, including the applicability of questions, for all articles evaluated by both members of the pair.

#### Article features

Region of origin was obtained for each article according to the corresponding author’s affiliation. In the two cases with two corresponding authors from different regions, we assigned the article to the region that had the most authors in the paper. Citations for all articles were obtained from Crossref on Oct. 10th 2019, using the rcrossref R package (Chamberlain et al., 2019).

Article size was defined in terms of number of labeled figure subpanels and tables in the main text, as we considered this to be more related to the amount of data presented in an article than text length. The presence of supplementary material and its size (similarly defined as the number of labeled figure subpanels and tables) were also collected. Preprints were further classified according to the position of their figures in the PDF file, which could be presented embedded in the text or separately in the end.

The subject area of preprints was obtained from bioRxiv based on the repository’s prespecified categories. In the only article listing two areas, the first one was considered. It was unavailable for one preprint. For PubMed articles in the independent sample, two researchers (C.F.C.D. and O.B.A.) independently assigned the article to one of the subject areas from bioRxiv’s classification. Disagreements were solved by discussion until consensus was reached. Articles that were not adequately described by any of the listed categories were classified as “other”. Peer-reviewed articles in the paired sample were assigned the same subject area as their preprint version.

#### Journal and Publisher Metrics

We obtained the impact factor for each journal according to the Journal Citation Reports from the corresponding year of online publication. Open-access status was attributed to journals listed on the Directory of Open Access Journals, assessed on Oct. 10th, 2019.

Journals were classified as “for-profit” or “non-profit” according to information obtained on their websites. “Non-profit” status was assigned to journals maintained solely by scientific societies or non-profit organizations. If a journal was associated with a scientific society but managed by a commercial publisher, it was classified as “for-profit”. From the journal’s or publisher’s online instructions to authors, we collected whether standard peer review was single-blind (reviewers’ identities are hidden, authors’ are known), double-blind (neither reviewers’ or authors’ identities are known during the process) or open (reviewers’ and authors’ identities are known to each other).

### Outcome measures and statistical analysis

#### Primary outcome

Our primary outcome was the comparison of overall reporting scores between the bioRxiv and PubMed groups (independent samples comparison) and between preprints and their peer-reviewed version (paired sample comparison).

#### Planned secondary outcomes

For the independent samples comparison, prespecified secondary outcomes included comparisons of general and subjective scores between bioRxiv and PubMed, and comparisons of general and specific scores between both groups for each biological model. Other planned secondary outcomes were correlations between the overall score with region of origin, article size and journal impact factor.

For the paired sample, prespecified secondary outcomes included comparisons of specific, general and subjective scores between preprints and peer-reviewed articles. Additionally, we planned comparisons of scores for each section of the questionnaire, comparisons of overall scores for each biological model, and correlations between overall and subjective scores. The difference in score between preprint and published version was used for planned correlations with article size, region of origin, journal impact factor, journal open access status, publisher commercial status and embedding of figures in the preprint version.

#### Exploratory analyses

All other outcomes presented were not preregistered and should be interpreted as exploratory. Moreover, as both samples partially overlap, it is important to note that some outcomes for the paired sample were planned after independent sample data had been analyzed in an exploratory manner. This was the case for the comparison of scores in individual sections of the questionnaire, as well as for the correlations between overall score with subjective scores, publisher commercial status and open access status of the journal, all of which were exploratory in the independent samples comparison.

Other analyses were exploratory in both stages of the study. These include (a) comparisons of reporting percentages for each question between groups, (b) correlations between overall score (for the independent samples comparison) or change in score (for the paired sample comparison) with subjective assessments and presence and size of supplementary material, and (c) correlation of overall, general, specific and individual section scores with study category. In the paired sample, we also correlated overall scores with whether a preprint had been published or not, time from preprint to peer-reviewed publication and number of citations.

For comparison of overall scores between preprints with and without embedded figures, we chose to aggregate data from both stages of the study in a single exploratory analysis in order to maximize sample size, as this comparison only included preprints. We also correlated subjective scores in both questions with embedding of figures.

Exploratory analyses to evaluate data consistency included (a) comparisons of mean evaluator agreement between preprints and peer-reviewed articles (combining both stages of the study), (b) assessment of evaluator bias by analyzing the interaction between individual reporting scores and evaluator identity and (c) correlations of overall scores from the same preprint in both stages of the study.

#### Statistical analysis

All comparisons between two groups were performed using Student’s t-test (for the independent samples comparison) or paired t-tests (for the paired sample comparison). Interactions between group and categorical variables (evaluator identity, biological model, region of origin and presence of supplementary material) were analyzed using 1- or 2-way ANOVA (with repeated measures in the paired sample comparison). Correlations between quantitative variables were assessed by Spearman’s (number of main and supplementary figures, impact factor and citations) or Pearson’s (scores from each stage of the study, time from preprint to peer-reviewed publication, subjective assessment) coefficients. Comparisons for reporting percentages of individual questions were performed using Fisher’s exact test or McNemar’s exact test for the independent and paired samples comparisons, respectively.

Differences in the primary outcomes were analyzed for significance using α=0.05. To account for multiple comparisons, we used Sidak’s α correction for the secondary outcomes in each of the two stages of the study (independent and paired sample comparisons). Significance thresholds were adjusted for 15 comparisons (αadjusted=0.003) and for 4 correlations (αadjusted=0.013) for secondary analyses in the independent sample, excluding the preregistered primary outcome and exploratory analyses. In the paired sample, they were adjusted for 26 comparisons (αadjusted=0.002) and for 4 correlations (αadjusted=0.013). We present unadjusted p values for all comparisons, but they should be interpreted according to the number of comparisons performed as described above. Although we also present p values for exploratory analyses, we refrain from labeling any of them as statistically significant, following best-practice recommendations for statistical analysis (Wasserstein & Lazar, 2016).

The complete dataset obtained is provided as **Supplementary File 1**. All analyses were performed using R (v. 3.5), and the analysis script is available as **Supplementary File 2**. Data is presented throughout the text as mean ± standard deviation. Lines in graphs always represent mean values.

### Sample size calculation

#### Independent sample comparison

Sample size was calculated to detect a difference of at least 10% between groups in the primary outcome with 90% power at α=0.05, based on the coefficients of variation for the reporting scores obtained from a blind pilot analysis of the first 10 articles in each group, which had mean values (± S.D.) of 67.9 ± 10.6 for PubMed and 65.0 ± 13.1 for bioRxiv. This yielded a sample size of 76 articles per group, with each evaluator analyzing between 25 and 32 articles in this stage.

#### Paired sample comparison

Sample size was calculated to detect a difference of at least 5% between groups in the primary outcome with 90% power at α=0.05. We chose this difference instead of 10% at this stage in order to be able to detect the effect size found in the independent samples comparison. In a blind pilot analysis of the first 10 pairs, we obtained a mean difference between pairs (± S.D.) of 6.04 ± 9.03, and this standard deviation was used in the calculation. This resulted in an estimated sample size of 37 pairs. By the time this estimate was obtained, however, we had already begun the evaluation of 56 pairs; thus, we decided to use this sample size in order not to discard any of the evaluations that had already been performed. With this sample size and the final S.D. for the difference between overall scores, our estimated power to detect a difference of at least 5% between groups was 99.1%.

## Results

### Evaluation of articles

The flowchart of screening and inclusion of articles can be visualized in **Figure 1**. Of the 76 preprints analyzed in the first stage, 49 had been published by Dec. 31st, 2018. Of these, 43 were included in the paired comparison. Additionally, 13 preprints that met inclusion criteria but were not included in the independent samples comparison (due to lack of articles in the same category in the PubMed sample) were included in the paired comparison. As these two stages were performed in different time periods, preprints included in both samples were fully reanalyzed by different trios of evaluators in the second stage to prevent time-related bias in analysis. As expected, there was a strong correlation between results for the same preprints in both stages (**Figure S1**; r=0.87, 95% C.I. [0.78, 0.93], p=1.97⨯10^−14^; Pearson’s correlation).

**Figure 1.**
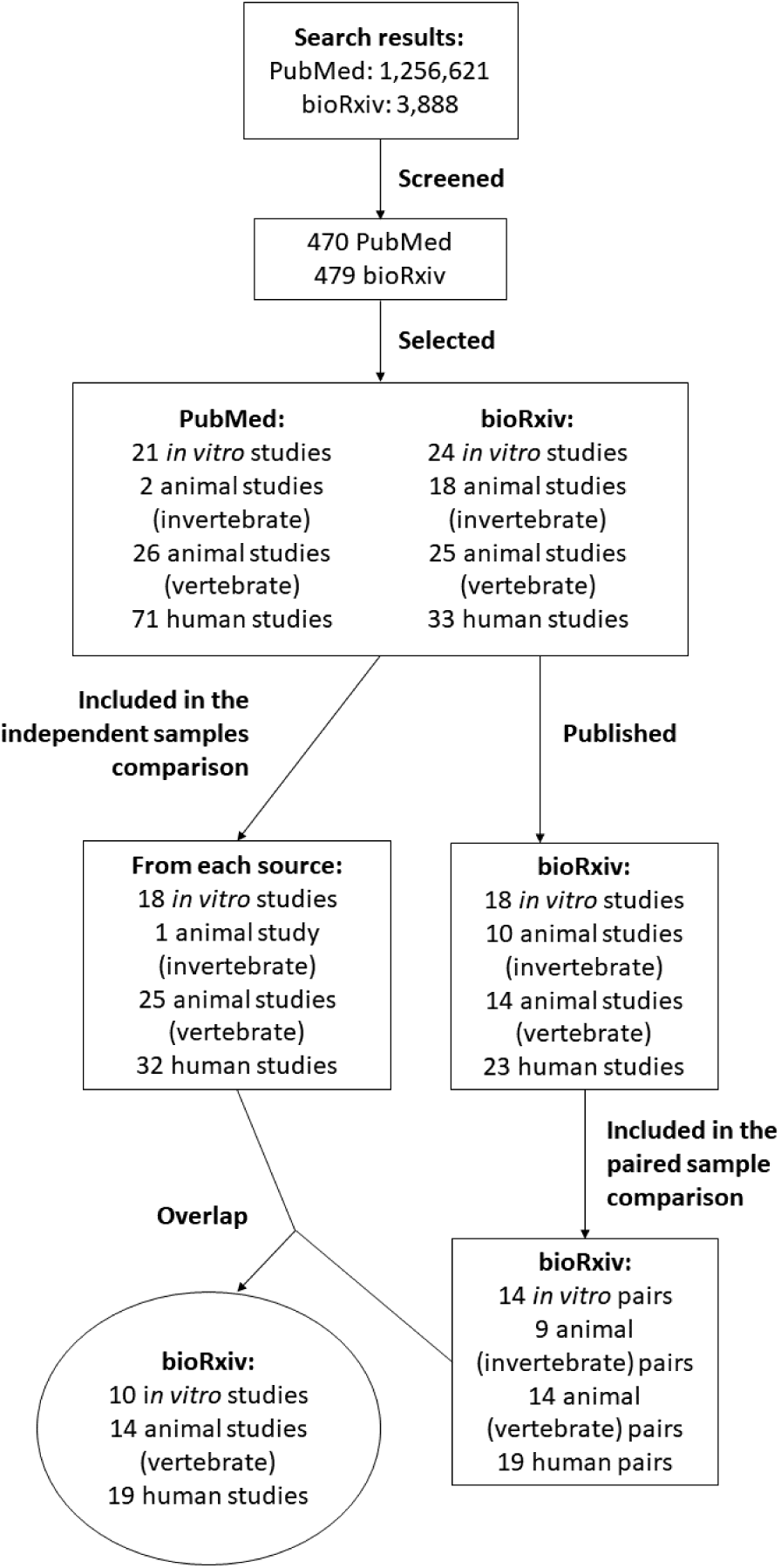
Flowchart of article screening and inclusion in each stage of the study. Screening and inclusion criteria are described in the Methods section.

17 out of 25 candidates reached criteria to be included as evaluators in the study. Two of them only participated in the independent samples comparison, while two others participated only in the paired sample stage. Agreement between evaluators after completion of data collection was above the test threshold for almost all evaluators (**Table S2**), with an overall agreement of 79.7%. There was no evidence of group bias by individual evaluators in either sample, as measured by interaction between evaluator identity and group in overall scores (**Table S3**; F=1.28, p_Interaction_=0.22 for the independent sample; F=1.05, p_Interaction_=0.40 for the paired sample; 2-way ANOVA). Mean agreement among evaluators was similar both in the independent samples comparison (81.1±6.8% vs. 79.3±5.9%, t=2.34, p=0.09; Student’s t-test) and in the paired sample comparison (78.4±7.1% vs. 78.3±7.4%, t=0.03, p=0.97; paired t-test).

### Article features

Adoption of preprints has been variable across different disciplines within the life sciences (Abdill and Blekhman, 2019; Sever et al., 2019). This can be clearly observed in our sample (**Table 1**), in which neuroscience articles account for almost half of bioRxiv articles included in the independent samples comparison, while prevalent areas in the PubMed group, such as clinical sciences and pharmacology, are underrepresented among preprints. There are also regional differences, with preprints more commonly coming from North America and Europe than PubMed articles (**Table 1**). The majority of vertebrate animal studies used rodents in both groups, although bioRxiv articles used mice more frequently than rats, while the opposite was seen in PubMed (**Table 1**). bioRxiv articles in the paired sample followed the same pattern as the independent samples comparison, with which it partially overlapped (**Table S4**).

**Table 1.** Sample description. Number of articles in each group by geographic region, main subject areas and animal species used. Only the most prevalent areas and animal models for both databases are shown; the complete data is available in **Table S4**.

| Region of origin | bioRxiv | PubMed |
| --- | --- | --- |
| North America | 34<br>44.7 % | 23<br>30.3 % |
| Europe | 32<br>42.1 % | 27<br>35.5 % |
| Asia | 5<br>6.6 % | 18<br>23.7 % |
| Other regions | 5<br>6.6 % | 8<br>10.5 % |
| Subject Area |  |  |
| Neuroscience | 34<br>44.7 % | 7<br>9.2 % |
| Pharmacology and Toxicology | 0<br>0 % | 12<br>15.8 % |
| Clinical Trials | 0<br>0 % | 9<br>11.8 % |
| Epidemiology | 0<br>0 % | 9<br>11.8 % |
| Microbiology | 7<br>9.2 % | 5<br>6.6 % |
| Physiology | 1<br>1.3 % | 6<br>7.9 % |
| Genetics | 5<br>6.6 % | 1<br>1.3 % |
| Cell Biology | 6<br>7.9 % | 2<br>2.6 % |
| Other | 23<br>30.3 % | 25<br>32.9 % |
| Species (vertebrates) |  |  |
| Mice | 14<br>56 % | 7<br>28 % |
| Rat | 1<br>4 % | 12<br>48 % |
| Macaca sp. | 3<br>12 % | 0<br>0 % |
| Zebrafish | 3<br>12 % | 1<br>4 % |
| Other species | 4<br>16 % | 5<br>20 % |

### Overall reporting score

As defined in our preregistered protocols, the overall score comparison between preprints and peer-reviewed articles was the primary outcome in each stage of the study. When comparing bioRxiv and PubMed articles (**Figure 2A**), we found a small difference between scores favoring PubMed articles (5.0, 95% C.I. [1.4, 8.6]; t=2.75, p=0.007, Student’s t-test). When comparing preprints to their own peer-reviewed versions (**Figure 2B**), we found a similar difference favoring peer-reviewed articles (4.7, 95% C.I. [2.4, 7.0]; t=4.15, p=0.0001; paired t-test). While reporting improved on average from preprint to published version, 27% of pairs (15 out of 56) presented a decrease in reporting score. We then performed secondary analyses to inquire whether the differences observed could be explained by particular study features in each group.

**Figure 2.**
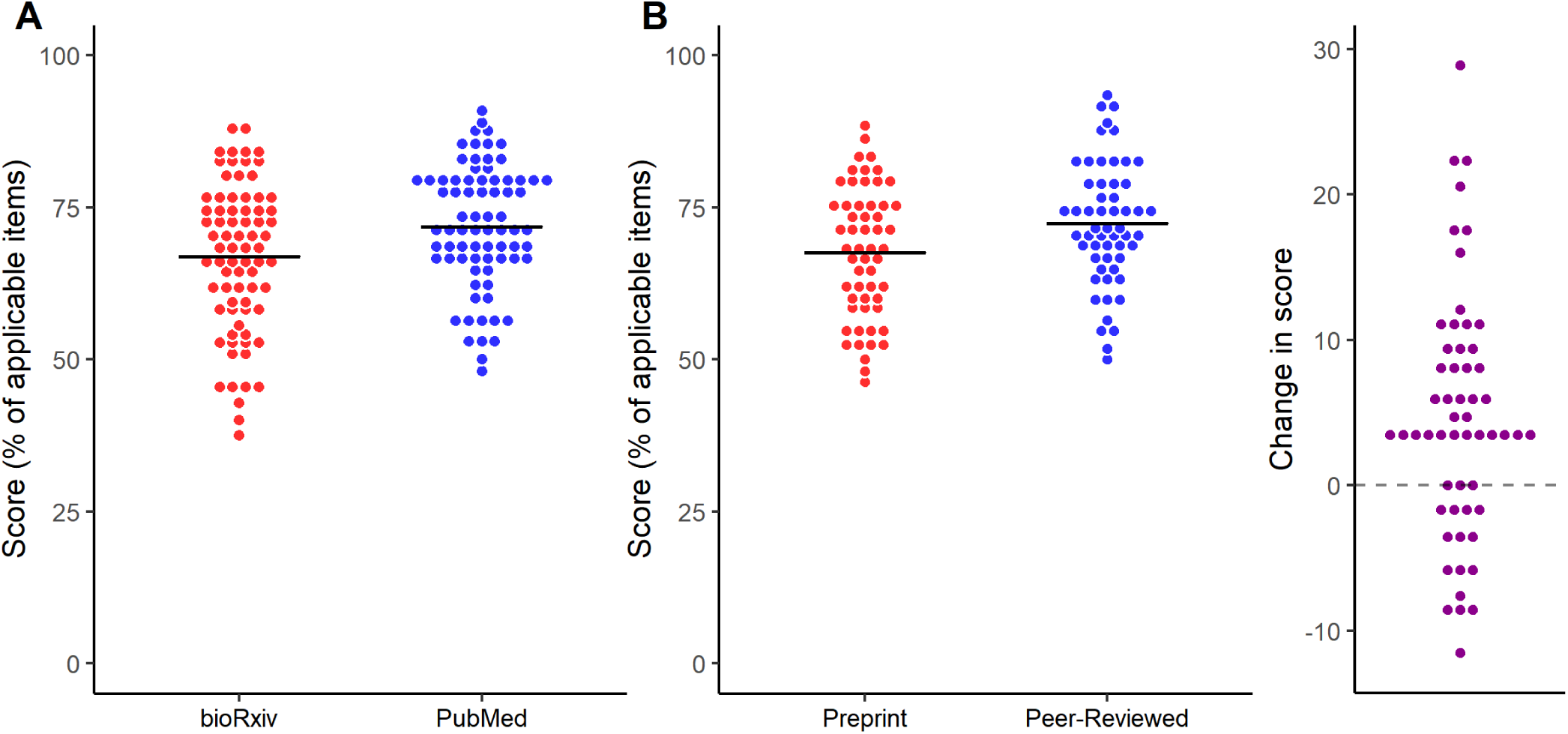
Reporting scores by source of the article. **(A)** Random samples of bioRxiv and PubMed were evaluated. Mean±S.D.: bioRxiv = 66.9±12.2, PubMed = 71.9±10.1; n=76/group. Student’s t-test, t=2.75, p=0.007, 95% C.I. [1.40, 8.59]. **(B)** A sample of bioRxiv articles was compared against their peer-reviewed version, published by a journal. Mean±S.D.: Preprint = 67.6±10.8, Peer-Reviewed = 72.3±10.1; n=56 pairs. Paired t-test, t=4.15, p=0.0001, 95% C.I. [2.44, 6.99]. On the right, absolute changes in score from preprint to peer-reviewed versions are plotted for each pair.

### Reporting scores by category of articles

We compared overall reporting scores, as well as those for the general and specific parts of the questionnaire, for each article category (e.g. *in vitro*, invertebrate, vertebrate and human studies). As shown on **Table S5**, the difference favoring PubMed articles was largely consistent across categories, both in the independent samples and paired sample comparisons. An exploratory 2-way ANOVA showed that article category had an important effect in reporting scores in all comparisons (a predictable finding, as the questionnaires themselves were different among categories); however, no evidence of interaction was observed for any of the comparisons (**Table S5**).

We then examined the individual sections that compose the general score (**Table 2**), to see whether differences between groups could be attributed to specific sections. In the independent samples comparison, the largest difference was found in the drugs and reagents section. This was also observed in the paired sample comparison, in which a large difference was also found in the risk of bias section. An exploratory interaction analysis shows that the difference between groups varied slightly according to the section of the questionnaire in both the independent and paired sample comparisons (p_Interaction_=0.04 for the independent sample, p_Interaction_=0.09 for the paired sample).

**Table 2.** Reporting scores by questionnaire section. Independent sample comparison p values are from Student’s t-tests, while paired samples’ ones are from paired t-tests. In the paired sample there were 32 preprints and 35 peer-reviewed articles with no applicable questions in the Drugs and Reagents section; thus, there were only 17 pairs available for the statistical comparison. 2-way ANOVA results are presented in individual lines below each comparison set. All values are presented as mean ± S.D. 95% C.I.; 95% confidence interval of the difference.

| Study stage | Subset | # of applicable questions | Score (preprints) | Score (peer-reviewed) | Difference [95% C.I.] | p value | Sample size |
| --- | --- | --- | --- | --- | --- | --- | --- |
| Independent samples | Title and abstract | 1 ± 0 | 84.2 ± 36.7 | 93.4 ± 25.0 | 9.2 [-0.8, 19.3] | 0.07 | 76 |
|  | Risk of bias | 3.7 ± 0.5 | 39.3 ± 23.6 | 39.9 ± 20.2 | 0.6 [-6.4, 7.6] | 0.87 | 76 |
|  | Drugs and reagents | 1.2 ± 1.4 | 62.2 ± 36.0 | 78.5 ± 29.2 | 16.3 [0.6, 31.9] | 0.04 | 31 (bioRxiv), 38 (PubMed) |
|  | Data presentation | 7.3 ± 0.9 | 75.1 ± 16.9 | 79.9 ± 12.7 | 4.8 [-0.02, 9.6] | 0.05 | 76 |
|  | Data analysis | 6 ± 0 | 83.6 ± 20.4 | 81.0 ± 19.4 | -2.6 [-8.9, 3.9] | 0.44 | 76 |
| Group: $F=5.53$, $df=1$, $p=0.02$; Section: $F=97.09$, $df=4$, $p<2\times 10^{-16}$; Interaction: $F=2.46$, $df=4$, $p=0.04$ | | | | | | | |
| Paired sample | Title and abstract | 1 ± 0 | 85.7 ± 35.3 | 83.9 ± 37.1 | 1.8 [-8.0, 4.4] | 0.57 | 56 |
| | Risk of bias | 3.7 ± 0.5 | 39.7 ± 20.6 | 54.2 ± 17.4 | 14.6 [8.4, 20.8] | $1.6\times 10^{-5}$ | 56 |
|  | Drugs and reagents | 0.9 ± 1.3 | 66.4 ± 35.7 | 81.6 ± 26.8 | 15.2 [4.0, 26.4] | 0.01 | 17 |
|  | Data presentation | 7.3 ± 0.8 | 78.4 ± 12.5 | 80.9 ± 13.3 | 2.5 [-0.3, 5.4] | 0.08 | 56 |
|  | Data analysis | 6 ± 0 | 84.5 ± 16.2 | 88.1 ± 14.2 | 3.6 [-0.5, 7.6] | 0.08 | 56 |
| Group: $F=6.31$, $df=1$, $p=0.01$; Section: $F=50.37$, $df=4$, $p<2\times 10^{-16}$; Interaction: $F=2.00$, $df=4$, $p=0.09$ | | | | | | | |

These observations are corroborated by an exploratory analysis of individual questions (**Table 3, Table S6**). In the independent sample, reporting of statements on conflict of interest (65.8% vs. 44.7%), presentation of a clearly defined variation or precision measure (84.5% vs. 66.2%), meaning of symbols used in figures (91.8% vs. 69.2%), supplier (88% vs. 48%) and randomization (47% vs. 0%) of experimental vertebrate animals and eligibility criteria of human subjects (90.6% vs. 59.4%) were higher in PubMed articles (p≤0.01, Fisher’s exact tests). Conversely, reporting of unit-level data was more frequently reported in bioRxiv articles (29% vs. 4.2%; p=4.5⨯10^−5^, Fisher’s exact test). However, the only question in which a clear difference in favor of peer-reviewed articles was observed in the paired sample comparison (**Table 3, Table S7**) was conflict of interest statement, although similar trends were observed in reporting of funding source and unit-level data. It should be noted that our sample size was planned for detecting aggregate differences; thus, statistical power for detecting differences in individual questions is rather limited.

**Table 3.** Frequency of reporting for individual questions. Only the questions with the largest differences in each comparison are shown; complete data is available in **Table S6** (for the independent samples) and **Table S7** (for the paired samples). As none of these items had articles scoring “Partially”, only the number and percentage of “Yes” answers are shown. Results are from Fisher’s exact tests (in the independent samples comparisons) and McNemar’s exact tests (in the paired comparisons). 95% C.I.; 95% confidence interval).

|  | Preprint | Peer-reviewed | Odds ratio<br>[95% C.I.] | p value |
| --- | --- | --- | --- | --- |
| **Independent samples** |  |  |  |  |
| Unit-level data | 22<br>29% | 3<br>4.2% | 0.1 [0.02, 0.4] | 4.5x10<sup>-5</sup> |
| Animal<br>source/supplier<br>(Vertebrate) | 12<br>48% | 22<br>88% | 7.6 [1.6, 49.9] | 0.005 |
| Eligibility criteria<br>(Human) | 19<br>59.4% | 29<br>90.6% | 6.4 [1.5, 39.8] | 0.008 |
| Randomization<br>(Vertebrate) | 0<br>0% | 8<br>47% | ∞ [2.0, ∞] | 0.003 |
| Symbol meaning | 27<br>69.2% | 45<br>91.8% | 4.9 [1.3, 23.0] | 0.01 |
| Conflict of interest<br>statement | 34<br>44.7% | 50<br>65.8% | 2.8 [1.2, 6.9] | 0.01 |
| **Paired sample** |  |  |  |  |
| Conflict of interest<br>statement | 26<br>46.4% | 47<br>83.9% | 11.5 [2.8, 100.6] | 1.9x10<sup>-5</sup> |
| Funding source | 50<br>89.3% | 55<br>98.2% | ∞ [0.9, ∞] | 0.06 |
| Unit-level data | 14<br>25% | 19<br>33.9% | ∞ [0.9, ∞] | 0.06 |

As conflict of interest statements are typically required during the submission process, it could be argued that the large change observed in this item is not due to peer review itself, but rather to requirements set in place during the submission process. Removing conflict of interest alone from the reporting score in the paired sample, the difference in reporting scores from preprint to peer-reviewed article decreased from 4.7 (95% C.I. [2.4, 7.0]) to 3.3 (95% C.I. [1.1, 5.5]). Smaller differences were found in other items potentially associated with the submission process to a peer-reviewed journal, but these were still among the largest observed. Namely, funding source was added in the peer-reviewed version in 5 (8.9%) pairs and ethical approval of vertebrate animal studies was added fully or partially in 4 pairs (30.8% of applicable pairs). Ethical approval of human studies was added in 2 pairs and removed in 1 (10.5% and 5.3% of applicable pairs, respectively); however, 15 preprints with human studies (79%) already had ethical approval reported, against only 7 (53.8%) vertebrate studies; thus, there was less room for improvement in this category.

### Correlations between region of origin and article size with reporting score

Region of origin was initially classified in 6 categories (Africa, Asia, Europe, Latin America, North America and Oceania); however, due to the small sample size in some regions, we combined Africa, Latin America and Oceania into a single category (Other) for analysis (**Figure S2A**). In the paired sample, we used the same classification categories, but predefined that regions with less than 10 occurrences would be combined into the “Other” category (**Figure S2B**). We did not find evidence for an effect of region on quality scores (F=0.69, df=3, p=0.56; 2-way ANOVA) or an interaction of region with group differences (F=2.64, df=3, p=0.05 for interaction, 2-way ANOVA) in the independent sample comparison. There was also no effect of region of origin on the change in scores from the preprint to the peer-reviewed version in the paired comparison (interaction: F=0.47, df=2, p=0.63; 2-way repeated-measures ANOVA).

To test whether differences in article length could account for group differences in reporting scores, we looked for a correlation between the number of subpanels and tables in articles and their reporting score in the independent sample (**Figure 3A**). We found a negative correlation for the aggregate of articles (ρ=-0.31, 95% C.I. [-0.46, -0.15], p=9.5⨯10^−5^; Spearman’s correlation), mostly driven by the correlation in the bioRxiv sample (ρ=-0.35, 95% C.I. [-0.56, -0.11], p=0.002), although a weaker negative correlation was also observed in the PubMed group (ρ=-0.22, 95% C.I. [-0.44, 0.0006], p=0.05). In the paired sample, we had planned to seek a correlation between the difference in scores and the difference in number of figures between preprints and peer-reviewed versions (**Figure 3B**; ρ=-0.07, 95% C.I. [-0.33, 0.17], p=0.59; Spearman’s correlation). However, article size varied only slightly from preprint to their respective peer-reviewed versions (with a mean change ± S.D. in number of subpanels/tables of 1.02±11.5 and a median of 0). We also performed an exploratory correlation between difference in reporting scores and the mean numbers of subpanels/tables between the preprint and peer-reviewed version, which showed a weak positive trend (**Figure 3C**; ρ=0.24, 95% C.I. [-0.05, 0.49], p=0.08; Spearman’s correlation).

**Figure 3.**
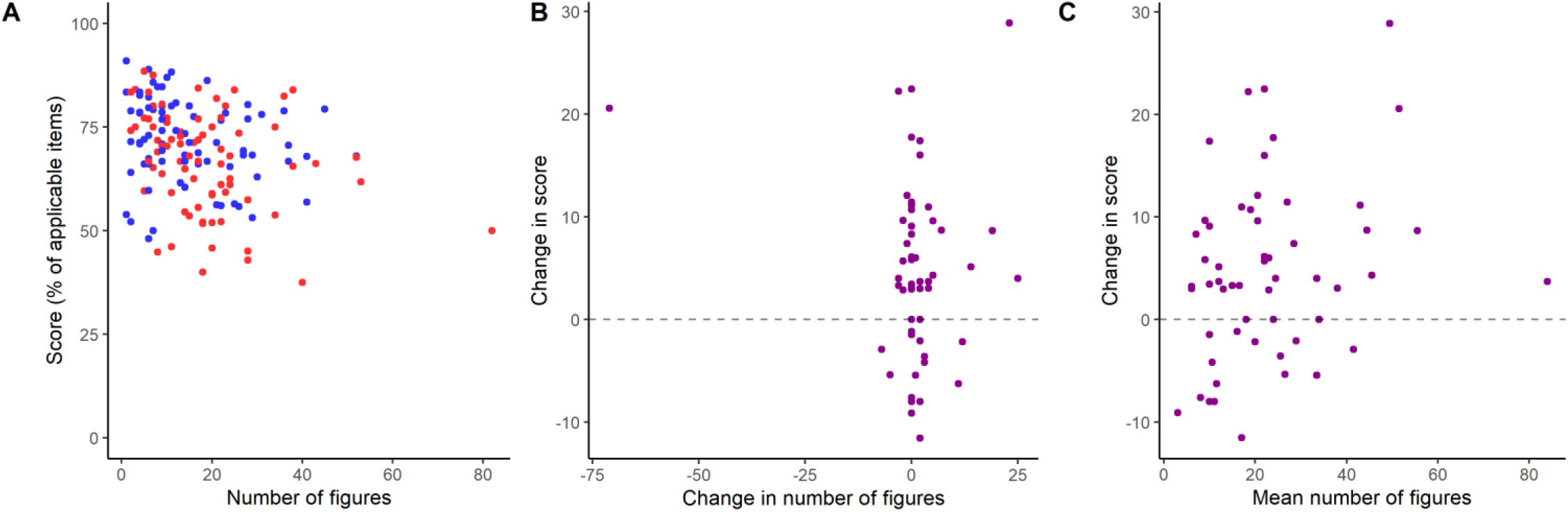
Quality of reporting by article size. **(A)** Overall scores by number of figure subpanels/tables in the independent samples comparison. Spearman’s correlations: All articles, ρ=-0.31, 95% C.I. [-0.46, -0.15], p=9.5⨯10^−5^; bioRxiv (shown in red), ρ=-0.35, 95% C.I. [-0.56, -0.11], p=0.002; PubMed (shown in blue), ρ=-0.22, 95%C.I. [-0.44, 0.0006], p=0.05. N=152 (76/group). **(B)** Change in score from preprint to peer-reviewed versions by change in the number of figures subpanels/tables in the paired sample. Spearman’s correlation: ρ=-0.07, 95% C.I. [-0.33, 0.17], p=0.59, N=56. One article presented a large decrease in number of figures (−71 figures subpanels/tables), as it was published as a brief communication. **(C)** Difference between scores from peer-reviewed to preprint version by mean number of figure subpanels/tables between preprint and peer-reviewed version in the paired sample. Spearman’s correlation: ρ=0.24, 95% C.I. [-0.05, 0.49], p=0.08, N=56.

Preprints contained supplementary data more frequently (39, vs. 20 PubMed articles) and had more supplementary subpanels on average (18.9±15.9 vs. 7.6±4.6, mean±S.D.; 95% C.I. [3.8, 18.8]; Student’s t-test, t=3.01, p=0.004) than randomly selected peer-reviewed articles, and peer-reviewed versions in the paired sample had an average of 3.87 figures/tables added (95% C.I. [-0.38, 8.13]; Paired t-test, t=1.82, p=0.07). As exploratory analyses, we tested for correlations between the presence of supplementary material with overall reporting scores in the independent samples comparison (**Figure S3A**) or with difference in scores in the paired sample comparison (**Figure S3B**). No interaction between reporting score and presence of supplementary material was found (independent samples: F=1.05, df=1, p=0.31 for interaction, 2-way ANOVA; paired sample: F=0.21, df=1, p=0.81, 1-way ANOVA). Also as exploratory analyses, we looked for correlations between number of supplementary figures and overall scores (independent sample comparison, **Figure S3C**) or differences in scores (paired sample comparison, **Figure S3D**). In the independent samples comparison, number of supplementary figures subpanels/tables showed a weak negative correlation trend with overall reporting scores (ρ=-0.21, 95% C.I. [-0.49, 0.05], p=0.11; Spearman’s correlation), while in the paired sample it correlated positively with increase in reporting scores (ρ=0.31, 95% C.I. [0.07, 0.52], p=0.02, n=56, Spearman’s correlation.

### Correlations between publication features and peer review with reporting scores

As publication venue is often (and controversially) used as a surrogate for quality assessments, we looked for a correlation of impact factor with reporting scores or changes in reporting score from preprint to peer-reviewed publications. Mean (±S.D.) impact factor for PubMed articles in the independent sample comparisons was 3.3±2.1, ranging from 0.456 to 14.9, with no correlation with overall reporting score (ρ=-0.11, 95% C.I. [-0.35, 0.13], p=0.35, Spearman’s correlation; **Figure 4A**). Impact factors in the paired sample were on average higher than randomly selected articles from PubMed (mean±S.D. = 7.2±5.6, ranging from 2.11 to 28). Once more, we found no evidence of correlation between impact factor of the publication venue and the difference in scores from preprint to peer-reviewed version (ρ=0.16, 95%C.I. [-0.13, 0.43], p=0.25, Spearman’s correlation; **Figure 4B**), suggesting that improvements in reporting by peer-review are not strongly related to this particular metric.

**Figure 4.**
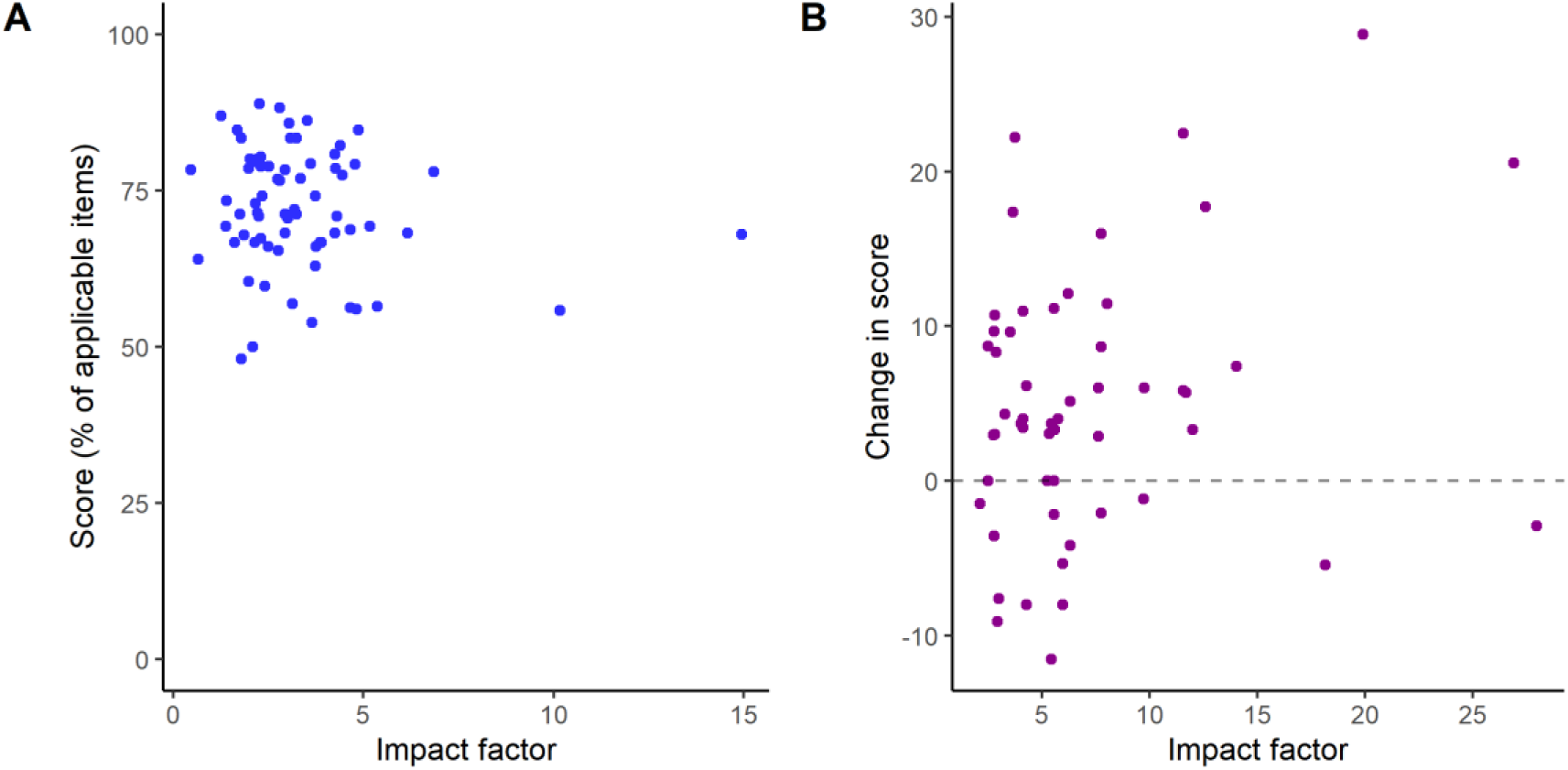
Quality of reporting by impact factor of publication venue. **(A)** Overall scores by 2016 impact factor of the publication venue in the independent samples comparison. Spearman’s correlation: ρ=-0.11, 95% C.I. [-0.35, 0.13], p=0.35, n=69. **(B)** Change in score from peer-reviewed to preprint version by impact factor of the peer-reviewed publication year in the paired sample. Spearman’s correlation: ρ=0.16, 95% C.I. [-0.13, 0.43], p=0.25, n=53. Impact factor was unavailable for 7 articles in the independent sample and 3 in the paired one.

We also looked for correlations between reporting quality and features of the publication venue, such as commercial and open-access status (**Figure S4A-D**). These analyses were exploratory for the independent samples comparison, but planned for the paired sample one. There was no correlation of commercial status of the publisher with reporting score (2.55, 95% C.I. [-3.59, 8.70]; t=0.83, p=0.41, Student’s t-test for the PubMed sample) or with changes in reporting scores (1.90, 95% C.I. [-2.78, 6.58]; t=0.81, p=0.42, Student’s t-test for the paired sample). Similarly, no correlation was found between open access status of the journal and reporting score (1.05, 95% C.I. [-4.52, 6.62]; t=0.37, p=0.71, Student’s t-test in independent sample) or changes in reporting scores (2.56, 95% C.I. [-1.99, 7.10]; t=-1.13, p=0.26, Student’s t-test in paired sample). It should be noted, however, that statistical power in these analyses was limited by the small number of journals in the open access and nonprofit categories.

In the paired sample, we also meant to explore correlations with features of the peer review process. However, the overwhelming majority of articles (36) were in journals that had single-blind review as the default option, while only 2 articles (from 2 different journals) had double-blind peer-review. Five articles are from journals in which authors choose between single- or double-blind peer review at submission, and three from journals in which authors choose between single-blind or open peer-review. Given the sample sizes, we decided not to perform any statistical comparisons.

We also collected the dates of submission to bioRxiv and dates of publication to assess whether the time lag between both – which might presumably correlate with the length of peer review in one or more journals – correlates with reporting quality. As observed in **Figure 5A**, there is considerable variation in time to publication (mean ± S.D. = 6.3 ± 4.1 months), and no correlation is observed with the change in score from preprint to published version (r=0.03, 95% C.I. [-0.23, 0.29], p=0.81; Pearson’s correlation). We also performed an exploratory comparison of reporting scores between preprints in the first stage of the study that had or had not been published by the end of 2018 (**Figure 5B**). Interestingly, we found a considerable difference, with preprints that were later published in a peer-reviewed journal having higher reporting scores on average (7.69, 95% C.I. [1.45, 13.93]; t=2.45, p=0.02, Student’s t-test), suggesting that reporting quality could have an impact on publication decisions by editors, reviewers, or by the authors themselves. Finally, we performed an exploratory correlation between number of citations and reporting scores in the independent samples (**Figure S4E**), with no correlation found in in either group (PubMed: ρ=-0.06, 95% C.I. [-0.27, 0.17], p=0.62; bioRxiv: ρ=0.10, 95% C.I. [-0.14, 0.32], p=0.38; Spearman’s correlation). We also found no clear correlation between total citations (sum of preprint and peer-reviewed versions) and changes in reporting scores in the paired sample(ρ=0.09, 95% C.I. [-0.19, 0.36], p=0.52; Spearman’s correlation; **Figure S4F**).

**Figure 5.**
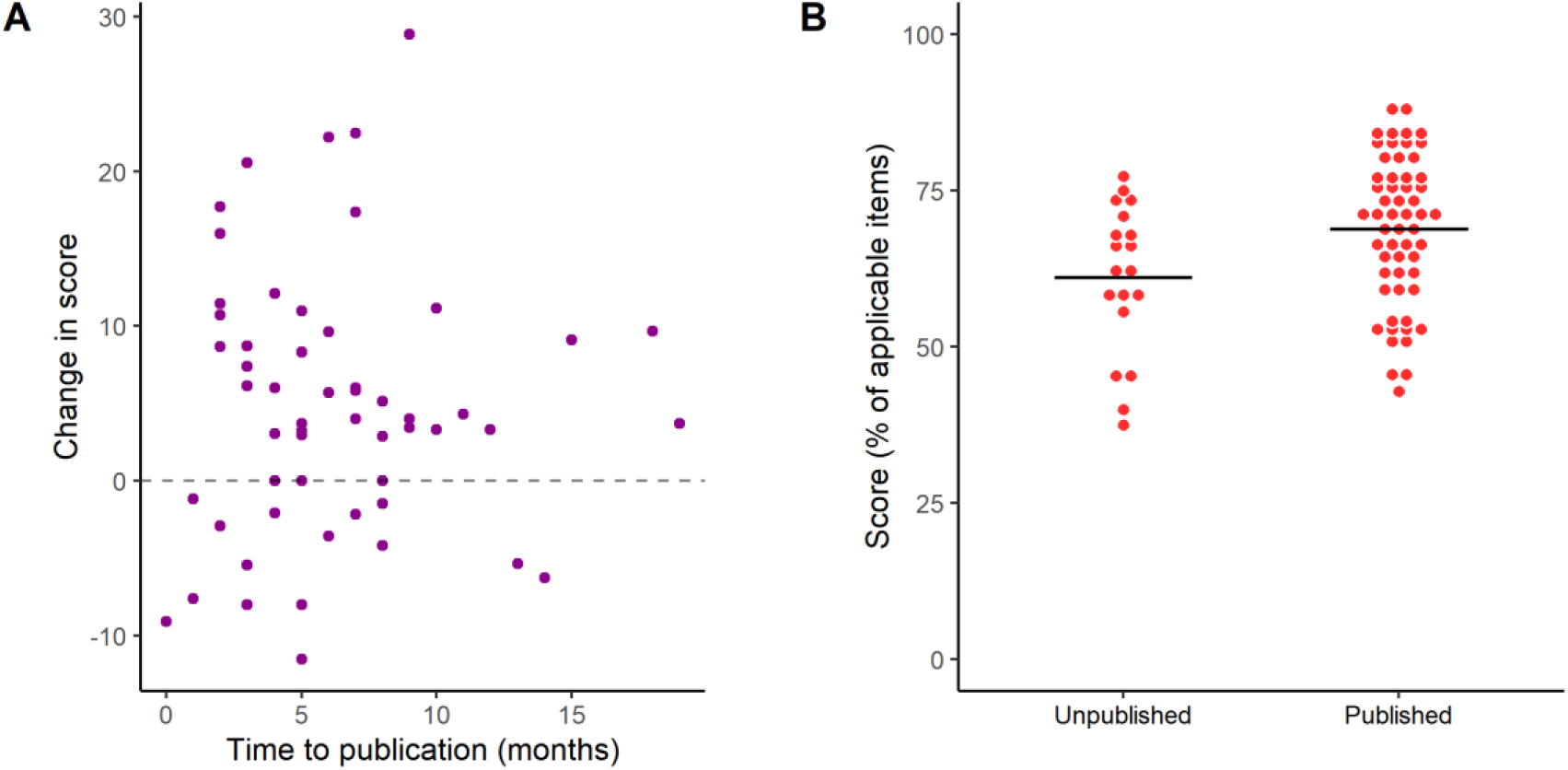
Quality of reporting and the peer review process. **(A)** Difference between scores from peer-reviewed to preprint version by time to publication (in months) in the paired sample. Pearson’s correlation: r=0.03, 95% C.I. [-0.23, 0.29], p=0.81, n=56 pairs. **(B)** Overall reporting scores by publication status (published or not in a peer-reviewed journal) of preprints assessed in the independent sample. Mean±S.D.: Unpublished = 61.1±11.9, n=19; Published = 68.8±11.8, n=57. 95% C.I. [1.45, 13.93], Student’s t test: t=2.45, p=0.02.

### Subjective assessment

As described in the Methods section, evaluators answered two subjective questions concerning the clarity of the title and abstract and the easiness to extract reporting information for the questionnaire. For clarity of abstract, we found a difference of 0.4, 95% C.I. [0.2 – 0.6] (t=3.61, p=4.2⨯10^−3^, Student’s t-test) in a 5-point scale favoring PubMed articles in the independent samples comparison (**Figure 6A**). In the paired sample comparison (**Figure 6B**), however, this difference was much smaller (0.2, 95% C.I. [-0.03, 0.4]; t=1.66, p=0.10, paired t-test), suggesting that difference between the PubMed and bioRxiv samples in abstract clarity is partially due to factors unrelated to peer review, such as subject area. Regarding easiness to extract information, there was again a large difference favoring PubMed articles in the independent samples comparison (**Figure 6C**; 0.7, 95% C.I. [0.5, 1.0]; t=6.22, p=5.1⨯10^−9^, Student’s t-test). This difference was also present, but smaller, when comparing preprints to their published versions (**Figure 6D**; 0.4, 95% C.I. [0.2, 0.6]; t=4.11, p=1.3⨯10^−3^, paired t-test).

**Figure 6.**
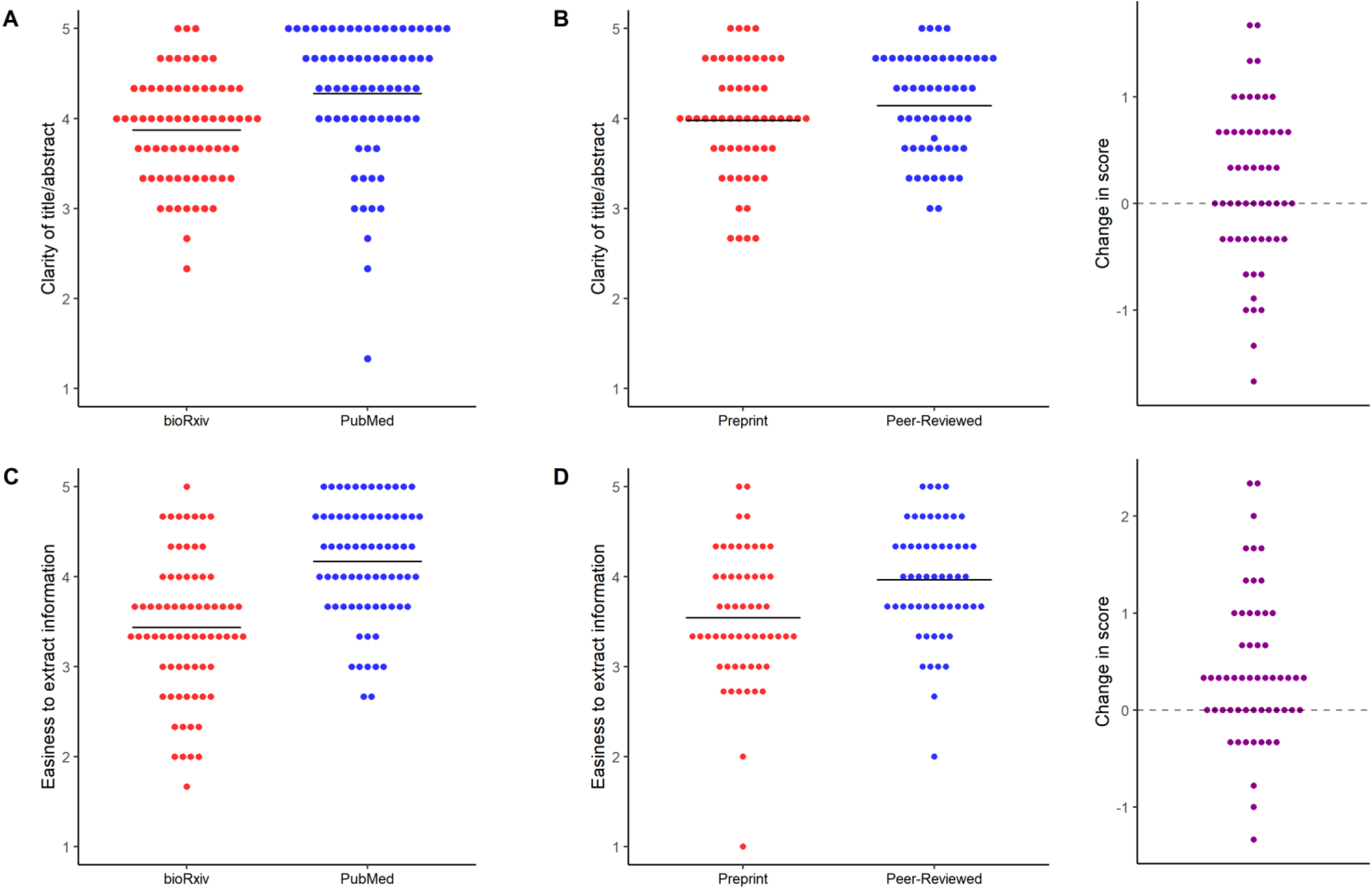
Subjective assessment by article source. **(A)** Clarity of title/abstract for the independent samples comparison. Scores were given as an answer to “Do the title and abstract provide a clear idea of the article’s main findings?”. Mean±S.D.: bioRxiv = 3.9±0.6, n=72; PubMed = 4.3±0.7, n=72. Student’s t-test: t=3.61, p=0.0004. **(B)** Clarity of title/abstract for the paired sample comparison. Mean±S.D.: Preprint = 4.0±0.6, n=56; Peer-reviewed = 4.1±0.5. Paired t-test: t=1.66, p=0.10. Right panel shows the differences between scores in preprint and peer-reviewed versions. **(C)** Easiness to extract information for the independent samples comparison. Scores were given as an answer to “Was the required information easy to find and extract from the article?”. Mean± S.D.: bioRxiv = 3.4±0.8, n=72; PubMed = 4.2±0.6. Student’s t-test: t=6.22, p=5.1⨯10^−9^. **(D)** Easiness to extract information for the paired sample comparison. Mean±S.D.: Preprint = 3.5±0.7, n=56; Peer-reviewed = 4.0±0.6, n=56. Paired t-test: t=4.12, p=0.0001. Right panel shows changes in score from preprint to peer-reviewed versions.

Based on the latter result, we questioned whether easiness to extract information could account for the difference observed in our primary outcome. To test this, we performed exploratory correlations between the two subjective questions and the overall reporting score for each group in the independent samples comparison, and with the change in score in the paired sample comparison. There was a strong correlation of clearness of title and abstract with reporting scores among PubMed articles (but not among bioRxiv ones) in the independent samples (**Figure 7A;** r=0.55, 95% C.I. [0.36, 0.69], p=6.3⨯10^−7^ and r=0.12, 95% C.I. [-0.11, 0.35], p=0.29 respectively; Pearson’s correlation); correlation between changes in this score and changes in reporting in the paired sample, however, was much weaker (**Figure 7B;** r=0.25, 95% C.I. [-0.01, 0.48], p=0.06, Pearson’s correlation). A strong correlation of easiness to extract information in both groups was also found with reporting scores in the independent samples comparison (**Figure 7C;** r=0.54, 95% C.I. [0.35, 0.68], p=1.04⨯10^−6^ for bioRxiv and r=0.60, 95% C.I. [0.43, 0.73], p=2.2⨯10^−8^ for PubMed), but the correlation between changes in subjective and reporting scores in the paired sample was again much weaker (**Figure 7D;** r=0.20, 95% C.I. [-0.06, 0.44], p=0.13, Pearson’s correlation).

**Figure 7.**
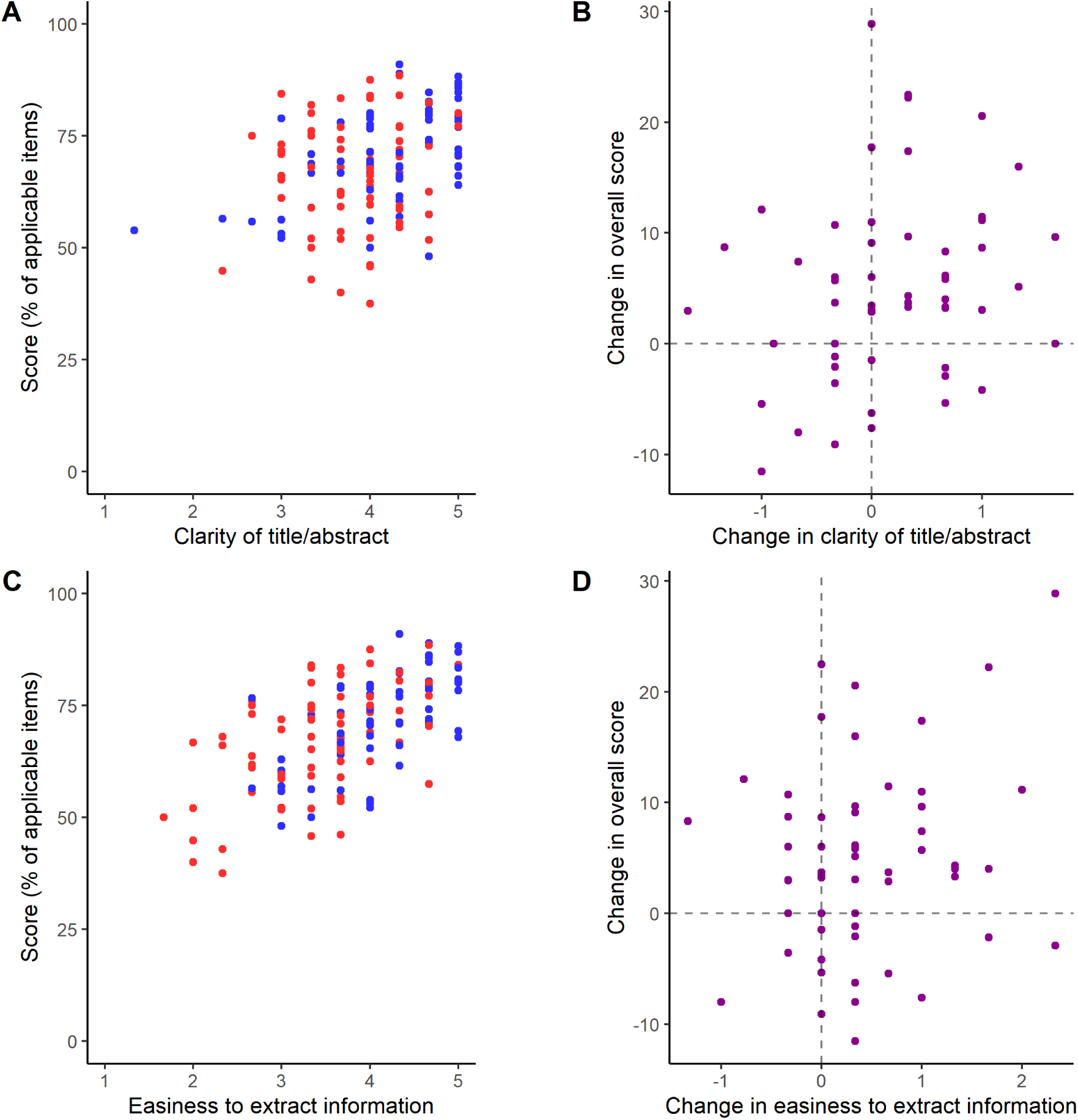
Quality of reporting by subjective scores. **(A)** Overall reporting scores by title/abstract clarity in the independent samples. Pearson’s correlation: r=0.38, 95%C.I. [0.23, 0.51], p=3.1⨯10^−6^, n=144 (all articles); r=0.12, 95% C.I. [-0.11, 0.35], p=0.29, n=72 (bioRxiv); r=0.55, 95% C.I. [0.36, 0.69], p=6.3⨯10^−7^, n=72 (PubMed). **(B)** Changes in overall reporting scores by changes in title/abstract clarity in the paired sample. Pearson’s correlation: r=0.25, 95% [-0.01, 0.68], p=0.06, n=56. **(C)** Overall reporting scores by easiness to extract information in the independent samples. Pearson’s correlation: r=0.59, 95% C.I. [0.47, 0.69], p=8.7⨯10^−15^, n=144 (all articles); r=0.54, 95% C.I. [0.35, 0.68], p=1.04⨯10^−6^, n=72 (bioRxiv); r=0.60, 95% C.I. [0.43, 0.73], p=2.2⨯10^−8^, n=72 (PubMed). **(D)** Changes in overall reporting scores changes in easiness to extract information in the paired sample. Pearson’s correlation: r=0.20, 95% C.I. [-0.06, 0.44], p=0.13, n=56. In all panels, bioRxiv articles are in red and PubMed ones are in blue, while differences between paired articles are shown in purple.

### Correlations between formatting and reporting score

Based on the correlation between easiness to extract information and reporting score, we inquired whether article formatting could influence both of these variables. As an exploratory way to assess this, we used our full sample of preprints (including those assessed in both the independent and paired samples stages) to compare those with figures at the end of the article to those with figures embedded in the text (which tend to be closer to the way data is presented in peer-reviewed articles) (**Figure 8A**). We found a small difference in reporting scores favoring the embedded group (70.8±11.6 vs. 64.6±10.9, Student’s t-test, t=2.37, p=0.02), which was similar in magnitude to that between the PubMed and bioRxiv groups. Both groups presented similar levels of improvement after peer-review in the paired sample (**Figure 8B**; 4.3±8.3 in the non-embedded group vs 5.6±9.0 in the embedded group; t=0.55, p=0.58, Student’s t-test). Nevertheless, there was no clear association of embedding with subjective assessments of title and abstract or easiness to extract information in the independent sample or with changes in these measures in the paired sample comparison (**Figure S5**).

**Figure 8.**
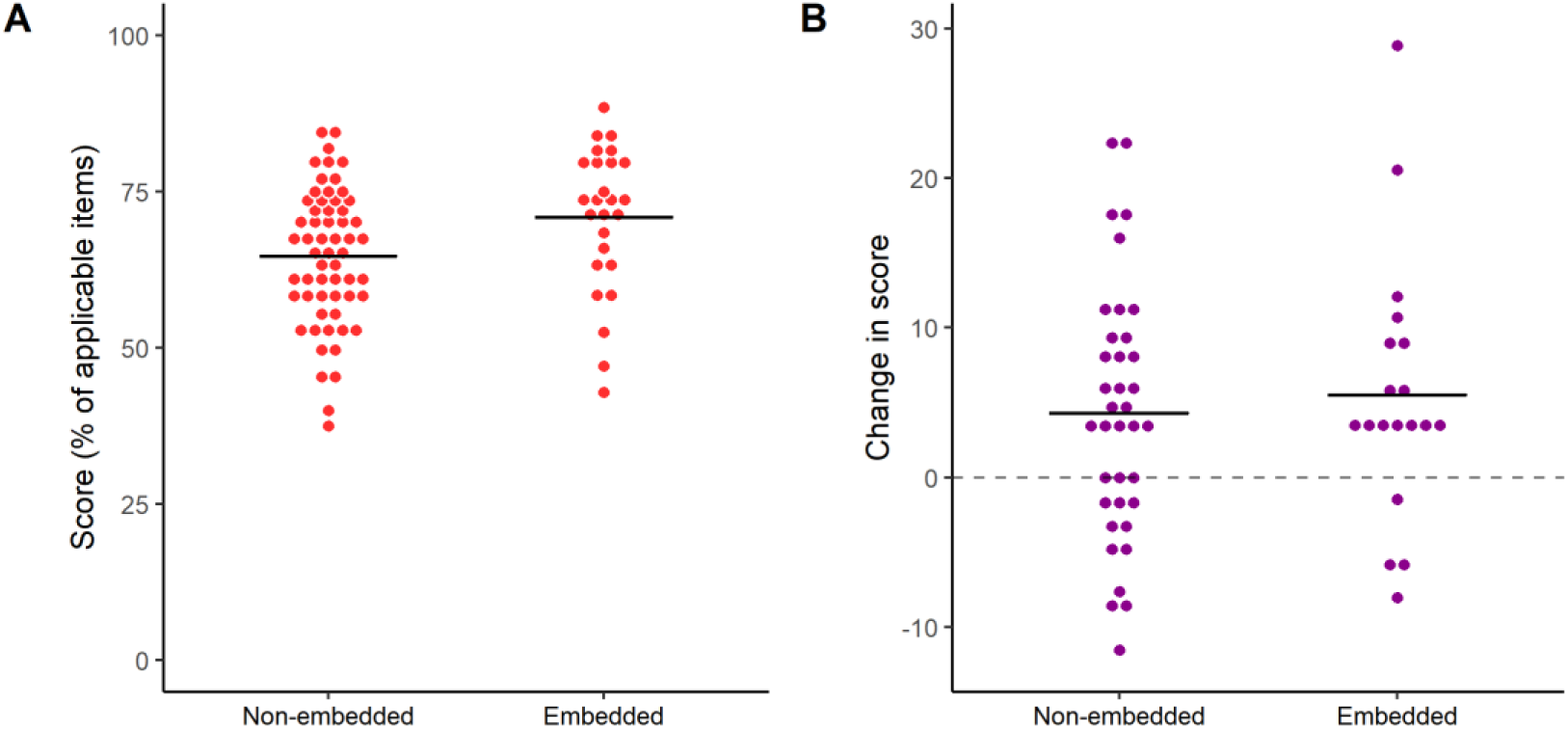
Reporting quality by article formatting. **(A)** Overall reporting score by embedding of figures in all preprints assessed. Preprints assessed in both stages of the study were included only once for this analysis, with the mean of reporting scores from both assessments. Mean±S.D.: Non-embedded = 64.6±10.9, n=59; Embedded = 70.8±11.6, n=26. 95% C.I. [1.0, 11.4]. Student’s t-test: t=2.37, p=0.02. **(B)** Difference between scores from peer-reviewed to preprint version by embedding of figures in the preprint version (paired sample). Mean±S.D.: Non-embedded = 4.3±8.3, n=38; Embedded = 5.6±9.0, n=18. 95% C.I. [-3.6, 6.1]. Student’s t-test, t=0.55, p=0.58.

## Discussion

Our study aimed to compare quality of reporting between preprints and peer-reviewed articles. Peer-reviewed articles had better reporting scores both when comparing independent samples from bioRxiv and PubMed and when comparing bioRxiv preprints to their own published versions. This difference was consistent across article categories and did not seem specific to any of them; however, it was small in magnitude, and variation ranges were largely similar between groups. Given the average number of applicable questions (26.6 and 25.6 in the independent and paired samples, respectively), the absolute differences of 5% and 4.7% observed in the independent and paired sample comparisons represent a difference in reporting of approximately 1 item.

The differences in the independent sample comparison could stem from many potential confounders. For example, there were large disparities in the scientific fields represented within each database; thus, the typical manuscript submitted to a PubMed journal may not be comparable to the typical article found on bioRxiv even before peer review occurs. Uptake of preprints by different communities within the life sciences has not been uniform (Anaya, 2016; Inglis and Sever, 2016; Abdill and Blekhman, 2019; Sever et al., 2019) and, although our sample does not exactly reflect bioRxiv’s distribution of subject areas (Abdill and Blekhman, 2019; Sever et al., 2019) because of the limitations imposed by our inclusion criteria, it was highly enriched in fields such as neuroscience and genetics. On the other hand, clinical research – which bioRxiv only started accepting in 2015 (Sever et al., 2019) – was infrequent among preprints, perhaps due to concerns over the potential ethical consequences of non-peer-reviewed material (Lauer et al., 2015; Tabor, 2016). The regional distribution of articles in both groups was also different, with a greater prevalence of articles from North America and Europe in the bioRxiv sample, but no evidence of interaction was observed between region of origin and reporting scores.

In the paired comparison, on the other hand, the differences observed are more likely related to the editorial process. Peer review likely accounts for some of the changes found, which were positive on average, but usually small. We also observed some small decreases in reporting scores after peer review in individual articles, although we cannot exclude that they are due to variability in evaluation, as preprints and their peer-reviewed versions were assessed independently. In any case, we failed to find interactions between changes from preprint to peer-reviewed version and features of the articles or journal of publication.

Nevertheless, peer review might not be the only factor affecting quality of reporting. In this respect, it is worth noting that the greatest difference observed from preprints to their peer-reviewed versions was the prevalence of conflict of interest statements, an item that is commonly required at journal submission. Thus, some of the observed changes could be attributable to other features of the editorial process, rather than to the actual feedback provided by reviewers. Additionally, as we looked only at the first preprint version, some changes could have derived from feedback received from other sources, such as comments on the preprint version or reviews from previous submissions.

Subjective ratings of how clearly titles and abstracts presented the main findings and how easy it was to locate relevant reporting information showed more robust differences favoring peer-reviewed articles, especially in the independent samples comparison. This could indicate that there are important differences between articles in both groups that were not assessed by our questionnaire, which focused on objectively measured reporting features. The fact that changes in subjective assessments did not correlate with changes in reporting score in the paired sample indeed suggests that they assess different dimensions of quality. It is also worth noting that the questionnaire was developed mostly with basic experimental research in mind; thus, information might be harder to find for articles with complex datasets in areas such as genomics, neuroimaging or electrophysiology, which were more frequently found in the bioRxiv sample. Similarly, evaluators from other areas of science might have had more difficulty interpreting titles and abstracts in these cases.

Even though we developed the questionnaire and manual to be as objective as possible, some items still required appropriate expertise or subjective assessment for correct interpretation. As most of our evaluators work in laboratory science, articles from other fields might have presented added difficulties. Although our high inter-rater agreement suggests that precision was reasonable, crowdsourced efforts such as these inevitably lead to heterogeneity between evaluators. On the positive side, they also dilute individual biases, a particular concern in our case, as evaluators were not blinded to the group of origin. Although blinding would have reduced risk of bias, it would also have required removing article formatting, which is arguably a contribution of the editorial process, and could have introduced errors in the process. Nevertheless, the homogeneity of the effect across different evaluators suggests that assessment bias was at most a minor issue.

While we looked at random samples from large platforms, the generalizability of our findings is limited by our inclusion criteria, which selected papers containing primary data and statistical comparisons, as well as by the use of a single preprint platform. Another limitation of our approach is that the use of the first table or figure for analysis meant that, especially in studies using human subjects, which typically start with a description of the study sample, the data under study were not always the main finding of the article. This might have been more common in larger articles with many datasets, as the number of figures correlated negatively with quality of reporting in both preprints and peer-reviewed articles. The limitations imposed by not selecting the main findings are mitigated when comparing the preprints to their own peer-reviewed publications, in which the data under study was the same in both versions; nevertheless, it could still be argued that the effects of peer review might have been different had we selected a central result in all cases.

Concerning formatting, the structure of preprints was more variable than that of peer-reviewed articles, as bioRxiv does not impose any particular style; thus, most preprints presented figures and/or legends separately from the description of results in the text. In an exploratory analysis of this variable, we found that preprints with embedded figures had a mean reporting score closer to that of PubMed articles (70.8±11.6 and 72.5±10.1, respectively). Although this comparison was observational and exploratory, with unbalanced sample sizes between groups, embedding figures within the text of preprints seems like a sensible and simple recommendation that could conceivably improve information retrieval from articles.

Previous studies comparing pre- and post-peer review manuscript in specific journals have found that the positive differences brought about by peer review were most evident in the results and discussion sections (Goodman et al., 1994; Pierie et al., 1996). In our independent samples comparison, we found that differences in overall score were attributable to better reporting of various individual items in PubMed articles, such as suppliers and randomization in animal studies and eligibility criteria in human studies; nevertheless, some data analysis issues were actually better reported on bioRxiv. As most of these differences were not present in the paired sample, we believe they are more likely due to differences in practices between the scientific fields represented in each sample, rather than actual effects of peer review.

The results of the paired sample comparison, on the other hand, suggest that editorial peer review itself has at best a small effect on quality of reporting. As described above, positive changes were mostly seen in items that might be automatically required by journals, such as conflict of interest statements and reporting of funding sources. Moreover, variables that could be associated with more rigorous quality assessment, such as journal impact factor and time to publication (which could correlate with longer reviews or multiple rounds of revision) did not correlate with changes in reporting. This does not exclude, of course, that larger peer review effects may exist on other facets of article quality: as orientations to reviewers are variable and typically nonspecific, the bulk of reviewers’ efforts might be focused on other issues. It does suggest, however, that quality of reporting is a largely overlooked feature during the peer review process.

A recent systematic review (Glonti, Cauchi, et al., 2019) analyzed descriptions of peer review in the scientific literature to identify tasks that reviewers were expected to perform. Assessment of adequacy to reporting guidelines were rarely mentioned, while other aspects of reporting – such as clarity of tables and figures and how data was collected – were more frequent. Most of the instructions to reviewers from medical journals in another study (Hirst and Altman, 2012) emphasized issues about general presentation, but varied a lot in how explicit and detailed they were. The depth of evaluation that editors expect from reviewers also varied, and was associated with some journal features, such as having professional or invited editors (Glonti, Boutron, et al., 2019).

Previous studies have also found that providing additional specialized review based on reporting guidelines led to small improvements to manuscripts, while suggesting reporting checklists to regular reviewers had no effect (Cobo et al., 2007, 2011). Reporting guidelines and checklists provided to authors during the review or manuscript preparation processes have been reported to cause modest improvements limited to a few items in in vivo animal studies (Han et al., 2017; Leung et al., 2018; Hair, Macleod and Sena, 2019; The NPQIP Collaborative group, 2019). Thus, the intuitive expectation that quality of reporting should be an aspect of study quality that is easily amenable to improvement by peer review does not seem to be confirmed by the available data.

It is interesting to note, nevertheless, that reporting scores were higher on preprints that were later published in peer reviewed journals than on preprints that had not been formally published within our time frame. This could indicate that the peer review process, even though it adds little in terms of quality of reporting, is effective as a filter and selects papers with better reporting for publication. However, this is a speculative interpretation, as we cannot be sure that preprints that were unpublished by the end date of our study were indeed submitted to a journal. Moreover, this comparison is also observational – thus, rather than influence the chances of publication itself, quality of reporting could be a proxy for other dimensions of quality that are more important in this process.

## Conclusions

In summary, our results suggest that quality of reporting among preprints posted in bioRxiv is within a comparable range to that of peer-reviewed articles in PubMed; nevertheless, there is on average a small difference favoring peer-reviewed articles. Our paired analysis provides evidence that the editorial process, which includes (but is not limited to) peer review, has positive but small effects on quality of reporting. Our results thus seem to support the validity of preprints as scientific contributions as a way to make science communication more agile, open and accessible. They also call into question the effectiveness of peer review in improving simple dimensions of research transparency, raising the issue of how this process could be optimized in order to achieve this more efficiently.

## Author contributions

C.F.D.C., T.C.M and O.B.A. designed the study. C.F.D.C. and V.G.S.Q. coordinated data collection and collected additional data. C.F.D.C., T.C.M, V.G.S.Q. and O.B.A. screened articles for inclusion. C.A.M.C., C.B.H., D.R., D.E.H., E.A.D-S., F.E., F.Z.B., G.D.G., I.R.C., K.L.H., L.v.E., M.M., P.B.T., R.J.A., S.J.B., S.F.S.G. and V.T.B. collected reporting data. C.F.D.C. and O.B.A. wrote the manuscript. All authors critically revised the manuscript and approved the final version.

## Acknowledgements

This manuscript is formatted based on the model of Finkelstein et al. available at https://github.com/finkelsteinlab/BioRxiv-Template.

