## Supplementary Text 1 for "Comparing quality of reporting between preprints and peer-reviewed articles in the biomedical literature"

**General instructions**

First, we'd like to thank you for your interest in participating in our project. As you might already know, our goal is to compare the quality of reporting in articles published as a preprint (i.e. not peer-reviewed) with those published in scientific journals indexed in PubMed (i.e. peer-reviewed articles). Our reporting quality questionnaire aims to be simple and quick to answer, based on common points drawn from reporting guidelines for different types of experimental studies.

This manual is meant to be a guide whenever you have doubts regarding what is to be considered when answering the questions. We tried to present some examples, but please do not restrict yourself by those. If there are still doubts, please contact us. General comments may also be expressed in the text box in the end of each session (in case you do so, please remember to specify the questions which you are referring to).

Alongside this manual, you will receive a list of the studies to be evaluated, with (a) pdf of the article and all supplementary material available, (b) the figure, table or subfigure which should be analysed and (c) the category of experiment (i.e. human studies, non-human animal studies (vertebrates or invertebrates) or *in vitro* studies). Note that some sessions contain questions referring to the study as a whole while some sessions contain questions referring only to the specific data under analysis. Only one of the sessions (session 3 – risk of bias) contains both questions on the whole study (questions 1-3) and on the specific data (question 4). Sessions 7-10 are to be answered only when they are applicable to the data, according to the category of experiment selected for analysis in the article. Note that throughout the form, you will be asked which is the figure or table under analysis and this must be fulfilled each time. Be sure to also check supplementary material whenever it is available for information not found on main text (e.g. details on methods). Remember to explain in the text box by the end of each session whenever you check "Partially" or "Not Applicable".

Link to the form: https://goo.gl/forms/h5WbCYlhWIghPdM12

**Session 1 – Identification**

1. "Respondent": your name.
2. "PDF name": enter here the pdf name of the article being analysed (corresponding to an identifier code containing one letter followed by numbers).
3. "Category of experiments": check the box that apply.
4. "Figure/Table": enter here the specific figure or table indicated for analysis.

**Session 2 – Title and abstract**

To answer this session, consider only information present in the title or abstract of the paper. Either one suffices: it is not necessary for the information to be present on both.

1. "Is the biological model / species of animal under study reported?": The biological model or species of animal used must be clearly interpretable (it is not necessary to include the official scientific nomenclature for species). For example, where "participants" or "volunteers" is stated it is clear that the title/abstract refers to human studies. However,

when a primary culture is mentioned without the species from which it was obtained, this is not sufficient to infer the biological model. If the title or abstract only refer to the biological model as “animals”, this is also not sufficient.

“Do the title and abstract provide a clear idea of the article's main findings?”: In this score you will evaluate your overall comprehension of the abstract; how well written it was to allow clear and easy understanding of the research problem, the methods used, and the results and conclusions obtained. It is not necessary to read the whole article to judge whether the abstract correctly summarizes the findings presented – on the contrary, the question is meant to judge whether reading the abstract alone can provide a good idea of what has been done.

**Session 3 – Risk of bias**

Questions 1-3 refer to the whole article. Question 4 refer to the data under analysis.

1. “Do the authors report their funding source(s)?”: This simply requires a statement describing the funding sources for the study; it is not required to describe the role of the funders.
2. “Is there a statement describing the presence or absence of conflict of interest?”: A statement is required to indicate either presence or absence of conflict of interest. Select the option accordingly.
3. “Is a sample size calculation reported?”: Sample size calculation could be just mentioned (without a full description of the methodology), presented with the parameters used for the calculation (e.g. effect size, variance, power), or not mentioned at all; please choose the corresponding option accordingly. The parameters for the calculation can be based for any outcome described in the article – it is not required that they pertain specifically to the data under study.
4. “Is assessment of outcome measures reported to be done in a blinded fashion?”: This refers to whether the person(s) assessing the specific data under analysis (e.g. behavioural assessment of animals, image analysis in a microscope) is blinded to the experimental groups (“Yes (blinded)”). This refer both to fully subjective analyses (e.g. behavioural assessment of subjects based on rating scales) and to partially automated analysis (e.g. computerized image analysis in which the experimenter must manually define a region of interest, for example). In cases in which outcome analysis is completely automated – i.e. there are no degrees of freedom that depend on the observer, or it is inherently not possible to blind the measurement, please check “Automated/Not applicable” and justify in the text box. Note that this question does not refer to group assignment or performing of experiments, only to assessment of outcomes.

**Session 4 – Drugs and reagents**

**From this session onward, only the figures/tables selected for analysis must be considered**. These questions refer only to experimental data, independent of the category of the experiment. If the data selected for analysis is observational, choose “Not Applicable” for these questions.

1. "Are the suppliers for drugs or other treatments in the data under analysis reported?": This question is applicable for any experiment involving pharmacological compounds or other treatments (such as diets) used as intervention. For treatments that are not commercially available (e.g. those prepared by the authors or collaborators), a description of the preparation procedure or reference describing it should be provided. Common reagents such as saline, DMSO, ethanol, among others, must be considered when answering this question if they are used as the intervention, but not if they are vehicle/control for the intervention. Note that for this question only supplier is required.
2. "Is every antibody used in the data under analysis linked to a citation, catalogue number, clone number or validation profile?": This question refers to antibodies used in any kind of experiment, either as a part of the intervention (e.g. function-blocking antibodies) or as part of the detection procedures (e.g. western blots). For all antibodies there must be either a catalogue number, a citation, a clone number or a validation profile. If no antibody is used, choose "Not applicable".
3. "For pharmacological interventions, is the dose/concentration reported?": This question is applicable only to pharmacological interventions, independent of the experiment category. The dose may be reported in any unit, as long as there is enough information to reproduce the experiment exactly. Consider pharmacological interventions all interventions in which a molecule or combination of molecules is administered to the biological model (drugs, siRNA, plants extracts, interventional diets, among others). If there is no pharmacological intervention, please choose "not applicable".
4. "For pharmacological interventions, is the vehicle reported?": This question is applicable only to pharmacological interventions in which the active ingredient (drug, siRNA, plant extract, food) is diluted or solubilized in a vehicle. If there is no pharmacological intervention, please choose "not applicable".

**Session 5 – Data presentation**

These questions should be answered for all categories of experiment. However, only the data selected for analysis must be considered.

1. "Are the groups compared clearly described?": This question refers to whether you can clearly assess the experimental intervention(s) or features defining each group. If a group is presented as a control, it must also be clearly described.
2. "Does the study provide a clear timeline for the experimental procedures or exposures and the measurement of outcomes in the data under analysis?": This may be present in figures or in the text. It is not required that the description is provided all at once, but the sequence of all experimental procedures performed in the biological model up to the point of tissue collection, including the time interval between them, must be present. Note that this information could be given together with other data present in the paper (in the methods section, for example) and not chosen for analysis in this study. In this case, it should still be considered. This question is not applicable for retrospective studies.
3. "Is a well-defined summary estimate (e.g. mean or median) of quantitative variables provided for each group? (If "Not applicable" is chosen, please provide the reason)": This could be described anywhere in the text (i.e. methods or results), figures or figure legends. It can refer to estimates given either in the figures, tables or text. The nature of the estimate (mean or median) must be clearly stated, and not merely inferred from

graphs. If “Not applicable” is chosen (for example, in the case of categorical variables), please provide the reason.

4. “Are findings presented with a well-defined measure of variation or precision (e.g. SD/SEM/X%CI)? (If “Not applicable” is chosen, please provide the reason)”: It must be clear what error bars in figures or “±…” in text represent, such as standard deviation, standard error of the mean or confidence intervals. For confidence intervals, the level of confidence must also be defined (e.g. 90%, 95%...).
5. “Are unit level data presented? (If “Not applicable” is chosen, please provide the reason)”: If individual results (for each experimental unit) are presented instead of or in addition to a summary estimate in a figure/table (e.g. in a scatter plot), check “Yes (in figures)”. If the paper presents the results as a summary estimate, but has an additional file containing unit-level data, check “Yes (raw data file)”. If both are available, check “Yes (both)”. If none of these are available, check “No”. In cases of categorical variables, check “Not applicable”. If “Not applicable” is chosen for any other reason, please provide the reason.
6. “Are all data shown in figures or tables clearly attributable to a specific experimental group/condition?”: It must be clear what each set of data in the graphs and figures represent, either through graphical representations (colours, symbols, patterns) or text (in the figure itself or in the legends).
7. “Are the units for each quantitative measure/indexes shown clearly described?”: Each variable measured must have units presented (usually in the figure axis or legend). In the case of indexes that do not have units, their calculation must be explained. In the case of categorical variables, check “Not applicable” and justify.
8. “Is the meaning of any symbols used in figures/tables (e.g. *, #, $^{a}$) clearly described?”: If the figure or table uses symbols or letters to depict statistical significance or other features, their meaning must be clearly stated. If no symbols are used, choose “Not Applicable”. Note that for this question you must not consider symbols identifying groups in the graph (e.g. in a scatter plot the control group is represented by triangles, while the treated are circles), as this will already been assessed in question 6.

**Session 6 – Data analysis**

These questions should be answered for all categories of experiment. However, only the data selected for analysis must be considered.

1. “Is the experimental unit used for analysis clear?”: This question refers to the experimental or observational unit in the samples compared by the statistical tests used (i.e. the unit in which sample size is described). Examples include patients, animals, experiments, cultures or replicates. Note that this question does not refer to whether the unit chosen is the most appropriate methodologically, only if it is clearly described.
2. “Is sample size reported for each group?”: Sample size could be reported as exact for each individual group (e.g. n = 13 (treated group), 11 (control)), or it could be given as range (e.g. n = 11-13) for multiple groups). Select the option accordingly. Sample size should be explicitly stated, and not merely inferred by the number of dots in a scatter plot, for example.
3. “Are the statistical tests used clearly described?”: The statistical test(s) must not only be cited but also clearly associated with the data under analysis. This could be mentioned anywhere in the text, figures, tables and legends.

4. "Are the variables and groups to which each statistical result refers to made clear?": It must be clear which data (e.g. groups, time points, variables) is being compared by each statistical test performed.
5. "Are the results of any statistical test in the figure (including omnibus and post-hoc comparisons) provided (as a p value or otherwise)?": All comparisons performed should be presented with their respective p-values. For example, if a two-way ANOVA is performed it yields three p-values (factor 1, factor 2 and interaction) – all of these must be present. If additional post-hoc analyses are performed, their p-value must also be present. If only part of the p values are presented (for example, with post-hoc results but not omnibus p value for ANOVA), please choose "partially".
6. "Are exact p values reported up to 2 decimal units (e.g. p=0.46, p=0.05, p<0.01)?": For this question to be answered as "yes", p values must be exact, in contrast to ranges such as p<0.05 or p>0.1, for example. The only exception is for values below 0.01 (which can be described as <0.01, <0.001, etc.).

**Session 7 – *In vitro* studies**

These questions should only be answered for data falling into the *In vitro* studies category. If that is not the case, please skip to the next session.

1. "Was the source of cell lines or microorganisms provided?": The text must describe how the microorganisms or cell lines were obtained, providing a reference if necessary. If it was obtained from a donation, the original source must be described (i.e. if it was obtained from ATCC or generated in the lab).
2. "For studies involving cell lines or microorganisms, do the authors report whether they have been authenticated recently (e.g., by STR profiling: within 1 year of use)?": For "yes", the text must explicitly describe that an authentication has been done and when it was performed. If date is not available or is more than 1 year before the experiments, please choose the corresponding option.
3. "Is the culture medium reported?": For this question, reporting of the growth medium allowing replication of the experiment must presented.
4. "Are culture conditions (temperature, [$CO_2$] and presence of $O_2$) reported?": This question is applicable for both microorganism cultures and cell lines. Temperature, $CO_2$ concentration and aerobic/anaerobic conditions must all be described at least for the control group.

**Session 8 – Animal Studies (invertebrates)**

These questions should only be answered for data in the category Animal studies (invertebrates). If that's not the case, please skip to the next session.

1. "Is the animal species reported? (if yes, please specify which species)": The species must be stated, but it is not necessary to include the official scientific nomenclature. If you can infer the species by the strain or genus mentioned in the article, you can consider species as reported. One must also specify whether animals were obtained from wild conditions or bred in a laboratory or other controlled environment.
2. "Is the strain of the animals reported?": The strain of the animals used must be clearly stated. Transgenic strains must also be considered. If there is no strain reported, please select "no" even if you believe the species used does not have multiple strains available – this information will be filtered in further analysis. This question is not applicable for wild animals.

3. “Is the sex of the animals reported?”: The sex of the animals must be clearly stated (male, female or mixed). If, due to methodological limitations, sex cannot be determined, choose “Not Applicable” and justify your answer (e.g. due to age, absence of phenotypical differences, etc.). If there is no sex reported, please select “no” even if you believe that this information is irrelevant to the species used.
4. “Is the age of the animals reported?”: Age could be given in days, weeks, months or years. Note that it can be given as an exact age or as a range: please select the option accordingly. The developmental stage of the animal (or embryo) can be considered as age range. In the case of wild animals, in which age may not be precisely obtained, please choose “Not Applicable”.
5. “Is the source/supplier of the animals reported?”: Authors must clearly describe where the animals were obtained (e.g. local colony, commercial supplier, wild animals, donation). In the case of wild animals, a brief description of the place of collection is enough.
6. “For *in vivo* pharmacological interventions, is the route of administration reported?”: This question is only applicable to pharmacological interventions. Consider pharmacological interventions by the same criteria described in session 4. Common routes of administration are oral administration, injections or dilution in the animal’s environment, but others may be applicable. If no pharmacological intervention was performed, please choose “not applicable”. This question is only applicable to *in vivo* interventions.
7. “If anaesthesia was performed, are type, route and dose/concentration described?”: This question is applicable for any experiment that uses anaesthesia, except when anaesthesia itself is used as an experimental intervention (in which case these questions will already have be answered in section 3) or when it is used as part of the euthanasia method (in which case it will be answered in the next question. All parameters mentioned must be described. This question is only applicable to *in vivo* interventions. If no anesthesia was performed (apart from the exceptions mentioned above), please choose “not applicable”.
8. “Is the method of euthanasia/tissue collection reported?”: The method of euthanasia or tissue collection must be described, including details on anaesthesia, if it was used as a part of the procedure. This question is only applicable if any intervention or analysis is done post-mortem or in a tissue removed from the live animal (both *in vivo* or *ex vivo* experiments); thus, it is not applicable for behavioural experiments, for example, in which all outcomes are assessed in live animals.

**Session 9 – Animal studies (vertebrates)**

These questions should only be answered for data in the category Animal studies (vertebrates). If that is not the case, please skip to the next session.

1. “Is the animal species reported? (if yes, please specify which species)”: The species must be stated, but it is not necessary to include the official scientific nomenclature. If you can infer the species by the strain and/or genus mentioned in the article, you can consider species as reported. One must also specify whether animals were obtained from wild conditions or bred in a laboratory or other controlled environment.
2. “Is the strain of the animals reported?”: The strain of the animals used must be clearly stated. Transgenic strains must also be considered. If there is no strain reported, please

select “no” even if you believe the species used does not have multiple strains available – this information will be filtered in further analysis. This question is not applicable for wild animals.

3. “Is the sex of the animals reported?”: The sex of the animals must be clearly stated (male, female or mixed). If, due to methodological limitations, sex cannot be determined, choose “Not Applicable” and justify your answer (e.g. due to age, absence of phenotypical differences, etc.). If there is no sex reported, please select “no” even if you believe that this information is irrelevant to the species used.
4. “Is the age of the animals reported?”: Age could be given in days, weeks, months or years. Note that it can be given as an exact age or as a range: please select the option accordingly. The developmental stage of the animal (or embryo) can be considered as age range. In the case of wild animals, in which age may not be precisely obtained, please choose “Not Applicable”.
5. “Is the number of animals housed together reported?”: The number of experimental animals being housed together (per cage, or other containers) should be clear, either as an exact number or as a range.
6. “Is the source/supplier of the animals reported?”: Authors must clearly describe where the animals were obtained (e.g. local colony, commercial supplier, wild animals, donation). In case of wild animals, a brief description of the place of collection is enough.
7. “Are animals reported to be randomized to experimental groups?”: This refers to whether random allocation of animals to different experimental groups was performed by experimenters. It is not necessary to describe the method of randomization. If it is not possible to randomize the experimental groups (e.g. cases in which the group is not actively determined by the experimenter, as in the case of transgenic animals, or comparisons between different species or sexes), select “Not Applicable” and explain. This question is only applicable to experiments using *in vivo* interventions.
8. “For *in vivo* pharmacological interventions, is the route of administration reported?”: This question is only applicable to pharmacological interventions in vertebrates. Consider pharmacological interventions by the same criteria described in session 4. Common routes of administration are oral administration, intravenous injection, subcutaneous administration and intraperitoneal injections, but others may be applicable. If no pharmacological intervention was performed, please choose “not applicable”. This question is only applicable to *in vivo* interventions.
9. “If anaesthesia was performed, are type, route and dose/concentration described?”: This question is applicable for any experiment that uses anaesthesia, except when anaesthesia itself is used as an experimental intervention (in which case these questions will already have be answered in section 3), or when it is used as part of the euthanasia method (in which case it will be answered in the next question). All parameters mentioned must be described. This question is only applicable to *in vivo* interventions. If no anesthesia was performed (apart from the exceptions mentioned above), please choose “not applicable”.
10. “Is the method of euthanasia/tissue collection reported?”: The method of euthanasia or tissue collection must be described, including details on anaesthesia, if it was used as a part of the procedure. This question is only applicable if any kind of analysis is done post-mortem or in a tissue removed from the live animal (both *in vivo* or *ex vivo* experiments); thus, it is not applicable for behavioural experiments, for example, in which all outcomes are assessed in live animals.

11. “Does the manuscript include an explicit statement of approval by a clearly identified ethics committee?”: The approval may or may not be clearly associated with a committee, choose the option accordingly. The inclusion of a protocol number is not necessary.
12. “Does the manuscript name the international, national or institutional guidelines followed?”: This question refers to whether, aside from receiving approval by an ethics committee, the experimental procedures and animal care followed an established animal welfare guideline (e.g. local institutional guidelines, the Guide for the Care and Use of Laboratory Animals from the NIH).

**Session 10 - Human studies**

These questions should only be answered for data in the Human studies category. If that’s not the case, please skip to the next session.

1. “Does the manuscript describe the recruitment process (including the target population)?”: Authors must describe how (e.g. recruitment advertising, medical consultations) and where (e.g. university, hospital, community) the recruitment process took place. In the case of retrospective studies, the process of selecting data and patients must be clearly described.
2. “Are the eligibility criteria adequately described?”: The criteria used when selecting and/or excluding participants that entered the recruitment process must be clearly described. Note that eligibility criteria are defined before the beginning of the experiment, and thus might not always correspond to the actual sample obtained.
3. “Is the sex of the subjects reported?”: Authors must describe how many participants from each sex are included in each experimental group. If only one sex was used in the study as described in the eligibility criteria (e.g. a study explicitly focusing in an exclusively male or female population), please choose “not applicable”.
4. “Is the age range of the subjects reported?”: The age range of the participants included in each group must be described, either as a full range or with a summary estimate combined with a measure of variance (e.g. standard deviation, interquartile interval, etc.). Note that this applies to the actual age range of the groups, and not to the age used as inclusion criteria of the study.
5. “Are subjects reported to be randomized to experimental groups?”: It is not necessary to describe the method of randomization. If it is not possible to randomize experimental groups (i.e. situations in which experimenters do not actively determine this, such as observational studies or comparisons between different sexes or ages), select “Not Applicable” and explain.
6. “Are the subjects reported to be blinded to the experimental group?”: This question refers to whether study subjects know which group they are assigned to (e.g. control or intervention/exposed). Note that blinding of the experimenters and those assessing outcomes has already been described in session 4. If this is not possible, please choose “not applicable” and justify.
7. “For pharmacological interventions, is the route of administration reported?”: This question is only applicable for pharmacological interventions. Consider pharmacological interventions by the same criteria as described in session 4. Common routes of administration are oral administration, intravenous injection and subcutaneous administration, but others may be applicable.

8. “Does the manuscript include an explicit statement of ethical approval and identify the committee(s) approving the study protocol?”: The statement of approval may or may not be presented clearly identifying the committee involved, select the corresponding option. It is not necessary to include a protocol number.
9. “Does the manuscript name the international, national or institutional guidelines followed?”: This question refers to whether, aside from getting an approval by an ethics committee, the experimental procedures were reported to have followed an established guideline (e.g. local institutional guidelines, NIH policy or the Declaration of Helsinki).
10. “Does the manuscript report that every subject signed an informed consent form?”: It must be clearly stated that every participant agreed to enter the experiment and signed an informed consent form prior to the beginning of the experiment. This question is not applicable for retrospective studies.

**Session 11 – Subjective assessment**

“Was the required information easy to find and extract from the article?”: In this score you will evaluate your overall subjective evaluation of the ease in which the information required in the previous sections was found in the article. This is meant to be a subjective assessment of the clarity of results reporting within the article.
