## Supplementary Tables and Figures for "Comparing quality of reporting between preprints and peer-reviewed articles in the biomedical literature"

**Supplemental Figures**

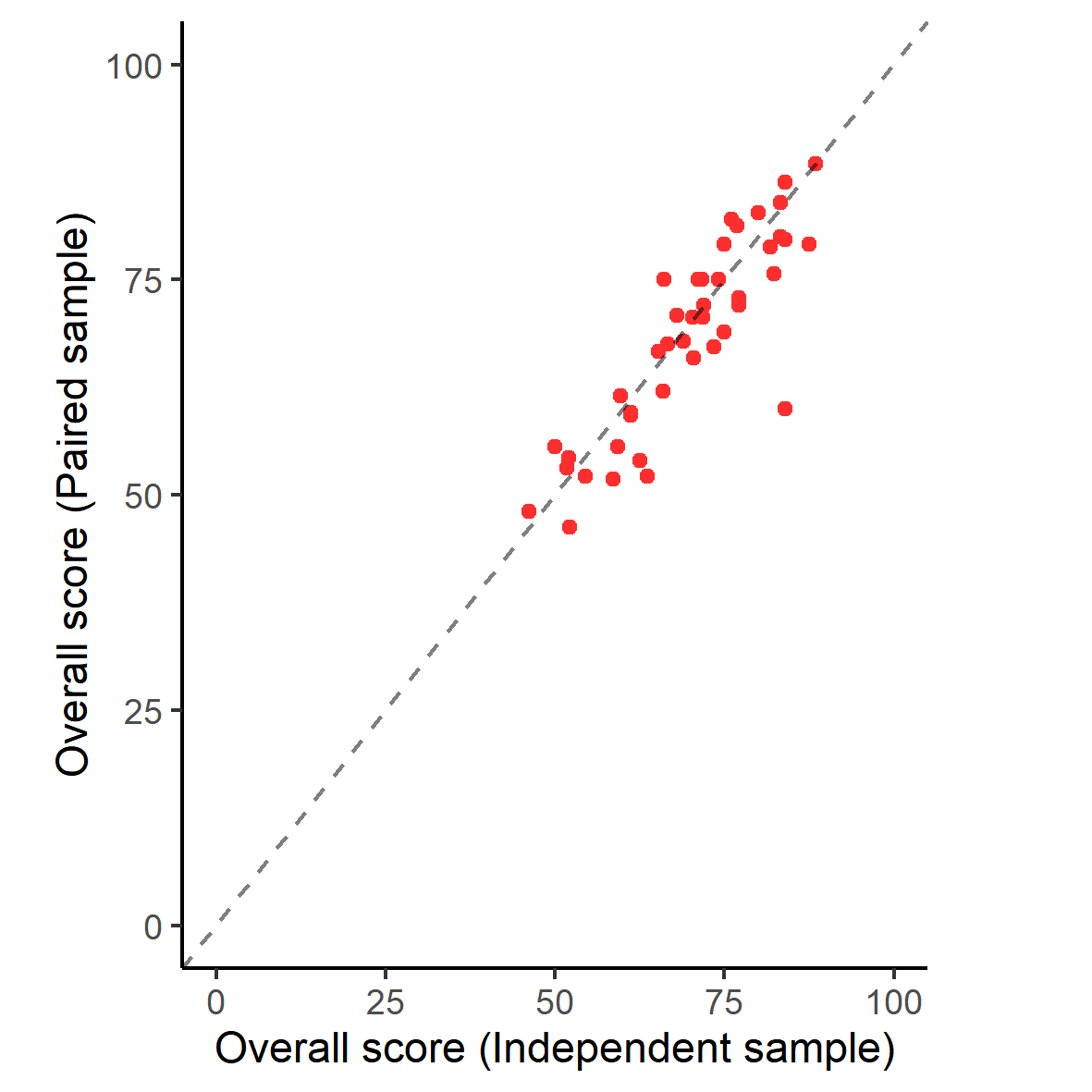

**Figure S1** – Correlation of reporting scores for the same preprints in the first (independent samples comparison) and second stage (paired sample comparison) of the study, in which they were analyzed by different evaluators. Pearson’s correlation: r=0.87, 95% C.I. [0.78, 0.93], p=1.97x10^-14^, n=43. Dashed line represents equality between both stages.

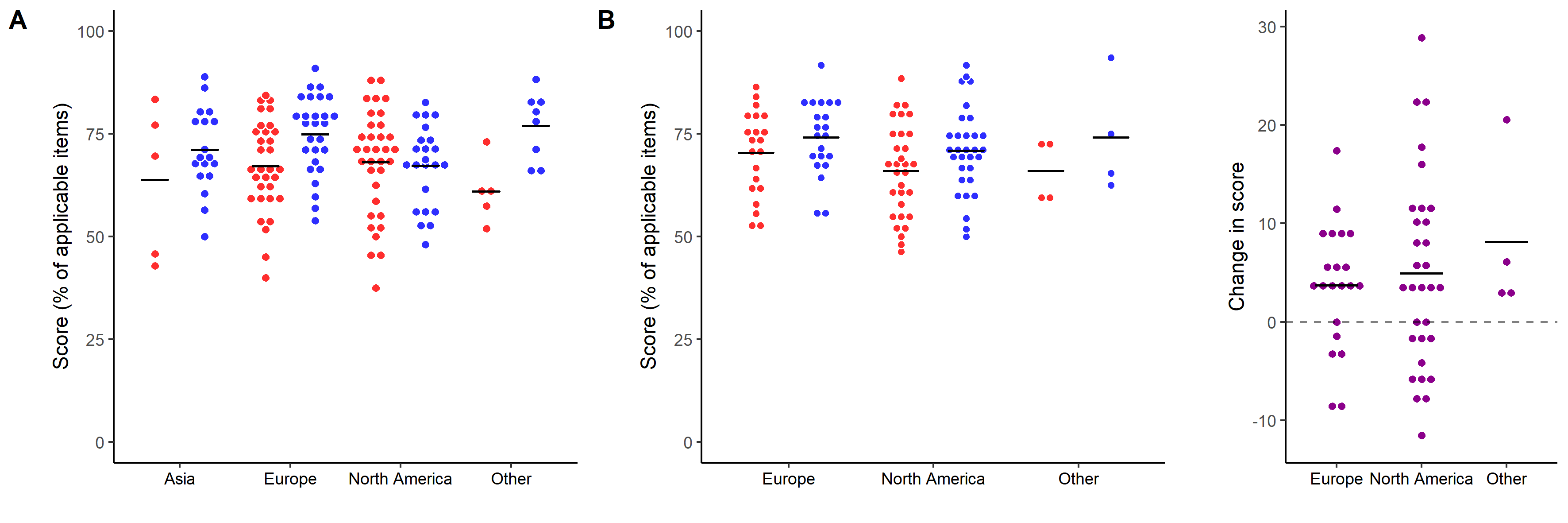

**Figure S2** – Quality of reporting by region of origin. **(A)** Overall reporting scores by region of corresponding author in the independent samples comparison. Two-way ANOVA: Group, F=7.75, df=1, p=0.006; Region, F=0.69, df=3, p=0.56; Interaction, F=2.64, df=3, p=0.05. bioRxiv articles are shown in red, while PubMed are shown in blue. **(B)** Overall scores by region of corresponding author in the paired sample. Two-way repeated measures ANOVA: Group, F=16.91, df=1, p=0.0001; Region, F=0.98, df=2, p=0.38; Interaction, F=0.47, df=3, p=0.63. Preprints are shown in red, while peer-reviewed articles are shown in blue. In the right panel, changes in score from preprint to peer-reviewed versions are plotted for each pair.

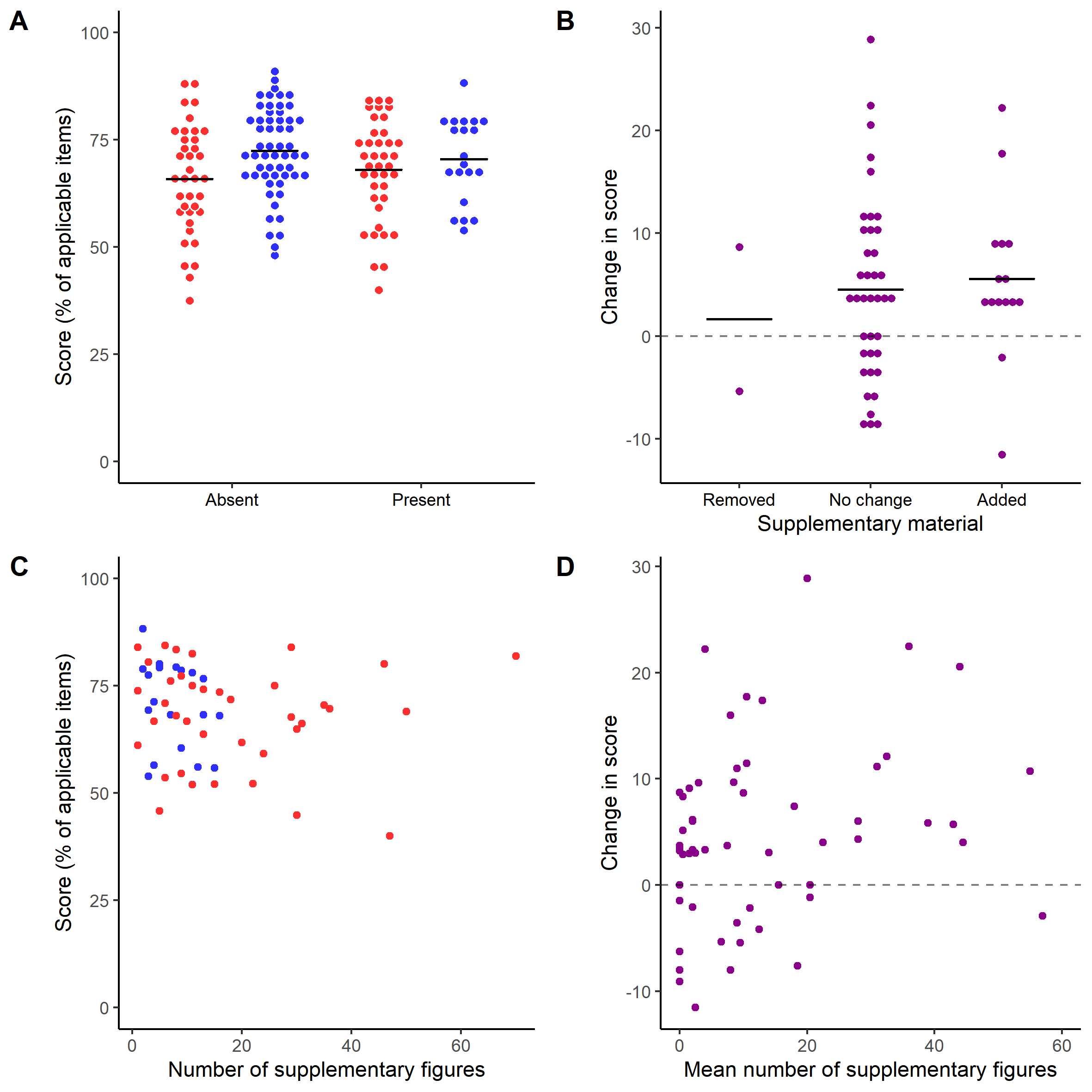

**Figure S3** – Quality of reporting by presence of supplementary material. **(A)** Overall reporting score by presence of supplementary material in the independent samples. Two-way ANOVA: Group, F=7.50, df=1, p=0.007; Suppl.Mat., F=0.04, df=1, p=0.85; Interaction, F=1.05, df=1, p=0.31. bioRxiv: n_Absent_=37, n_Present_=39; PubMed: n_Absent_=56, n_Present_=20. **(B)** Change in score from peer-reviewed to preprint version by change in supplementary material in the paired sample. One-way ANOVA, F=0.21, df=2, p=0.81. n_Removed_=2, n_NoChange_=39, n_Added_=15. **(C)** Overall reporting score by number of supplementary figure subpanels/tables in the independent samples. Spearman’s correlation: ρ=-0.21, 95% C.I. [-0.49, 0.05], p=0.11, n=59 (all articles with supplementary material); ρ=-0.13, 95% C.I. [-0.46, 0.23], p=0.42, n=39 (bioRxiv); ρ=-0.35, 95% C.I. [-0.74, 0.15], p=0.14, n=20 (PubMed). **(D)** Change in score from peer-reviewed to preprint version by mean number of supplementary figures. Spearman’s correlation: ρ=0.30, 95% C.I. [0.07, 0.52], p=0.02, n=56. In all panels, bioRxiv articles are in red and PubMed ones are in blue, while differences between paired articles are shown in purple.

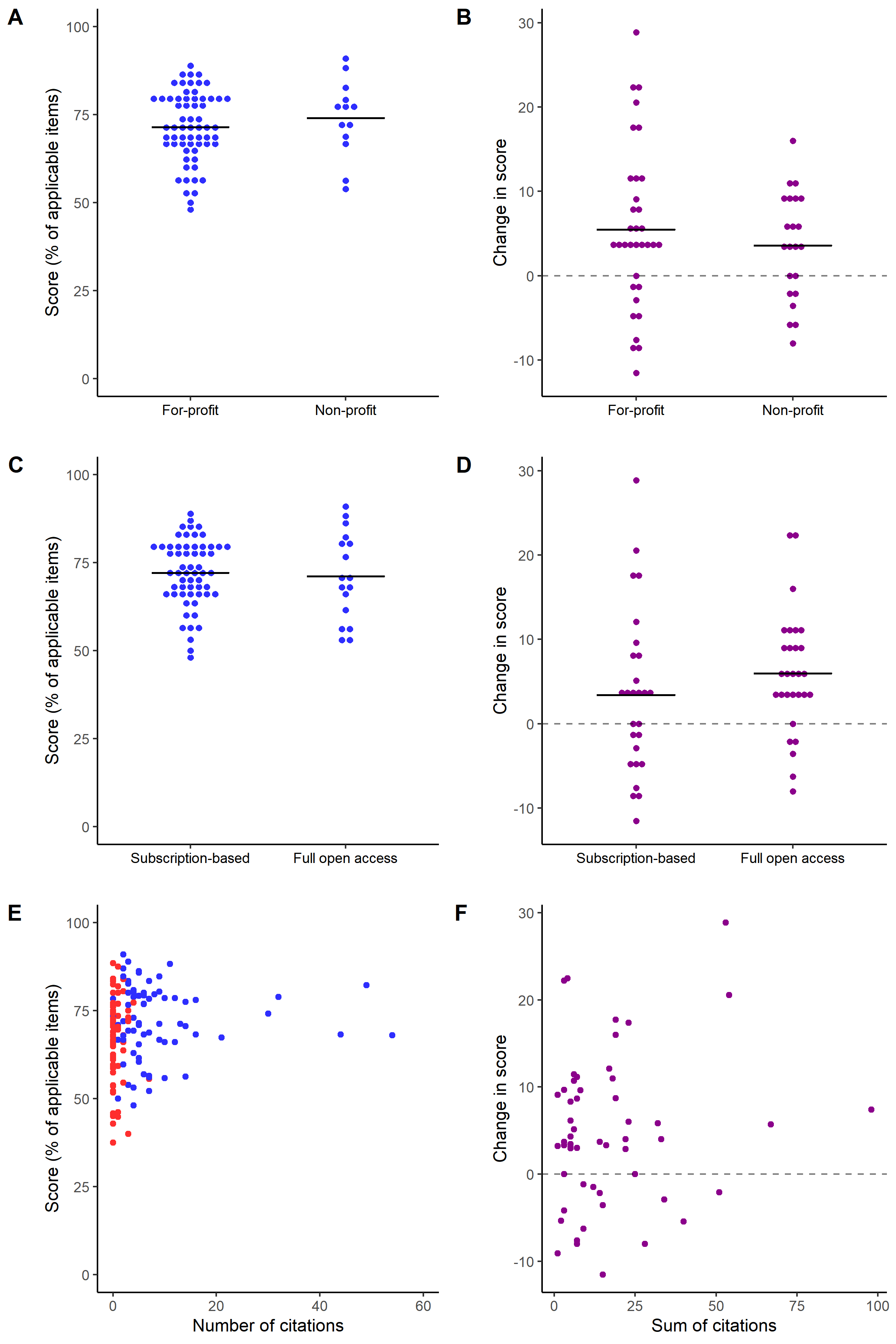

**Figure S4** – Quality of reporting by publication venue. **(A)** Overall reporting score by publisher classification in the independent sample. Difference, 95% C.I.: 2.5 [-3.6, 8.7]. Student’s t test, t=0.83, p=0.41. n_For-profit_=63, n_Non-profit_=13. **(B)** Change in score from peer-reviewed to preprint version by publisher classification in the paired sample. Difference, 95% C.I.: 1.9 [-2.8, 6.6]. Student’s t test, t=0.82, p=0.42. n_For-profit_=34, n_Non-profit_=22. **(C)** Overall reporting score by open access status of the journal in the independent sample. Difference, 95% C.I.: 1.0 [-4.5, 6.6]. Student’s t test: t=0.37, p=0.71. n_Subscription_=59, n_Open_=17. **(D)** Change in score from peer-reviewed to preprint version by open access status of the journal in the paired sample. Difference, 95% C.I.: 2.6 [-2.0, 7.1]. Student’s t test: t=-1.13, p=0.26. n_Subscription_=27, n_Open_=29. **(E)** Overall reporting score by number of citations in the independent sample. Spearman’s correlation: ρ=0.10, 95% C.I. [-0.14, 0.32], p=0.38, n=76 (bioRxiv); ρ=-0.06, 95% C.I. [-0.27, 0.17], p=0.62, n=73 (PubMed). **(F)** Change in score from peer-reviewed to preprint version by sum of citations in the paired sample. Spearman’s correlation: ρ=0.09, 95% C.I. [-0.19, 0.36], p=0.52, n=55. In all panels, bioRxiv articles are in red and PubMed ones are in blue, while difference between paired articles are shown in purple.

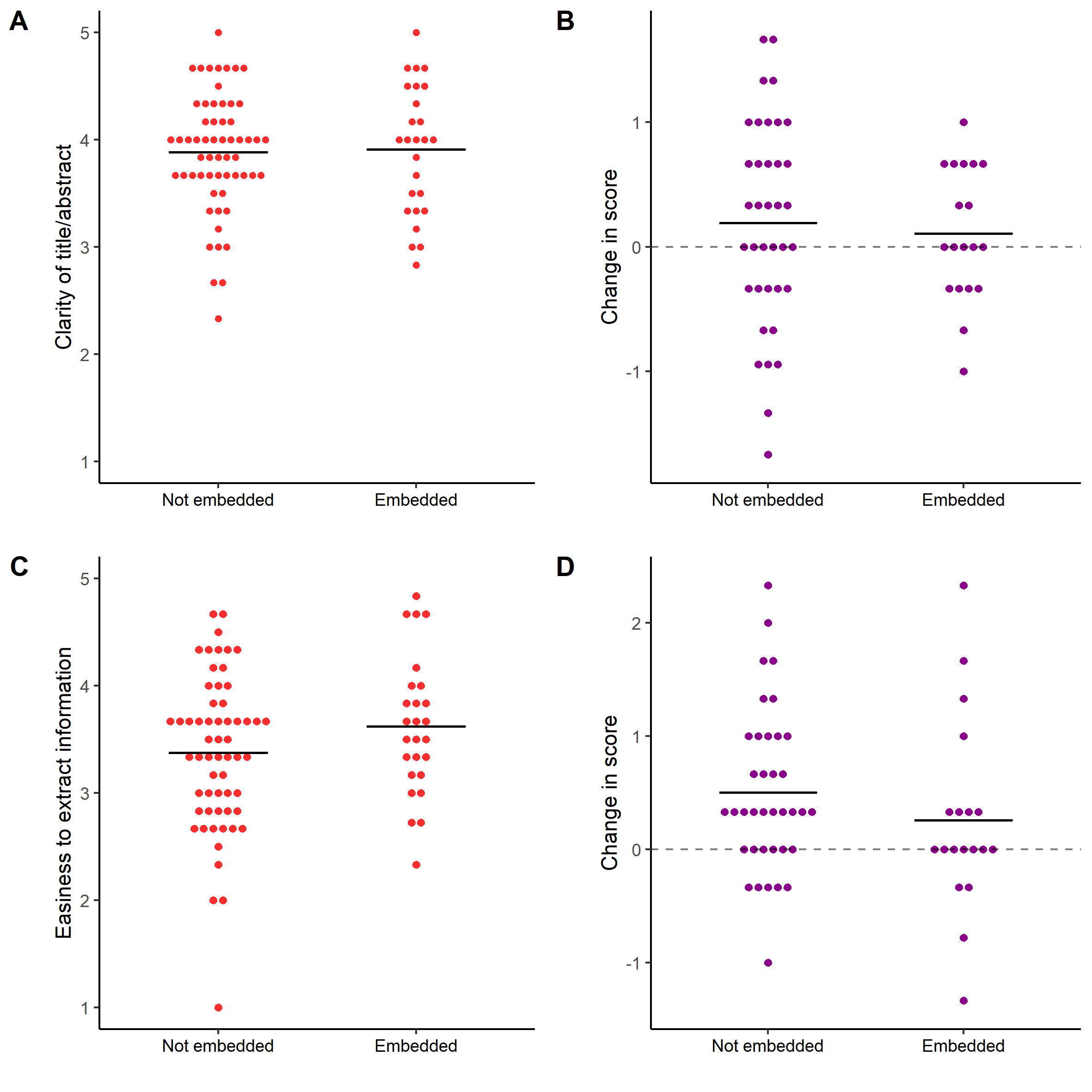

**Figure S5** – Subjective assessment by article formatting. **(A)** Mean subjective score for title/abstract clarity by embedding of figures in all preprints assessed. Preprints assessed in both stages of the study were included only once for this analysis, with the mean of subjective scores from both assessments. Difference, 95% C.I.: 0.03 [-0.23, 0.29]. Student’s t test: t=0.19, p=0.84 (n_Not_=59, n_Embedded_=26). **(B)** Difference between subjective scores for title/abstract clarity from peer-reviewed to preprint version by embedding of figures in preprints from the paired sample. Difference, 95% C.I.: 0.09 [-0.3, 0.5]. Student’s t test: t=0.36, p=0.72 (n_Not_=38, n_Embedded_=18). **(C)** Mean subjective score for easiness to extract information by embedding of figures in all preprints assessed. Preprints assessed in both stages of the study were included only once for this analysis, with the mean of subjective scores from both assessments. Difference, 95% C.I.: 0.2 [-0.08, 0.6]. Student’s t test: t=1.51, p=0.13 (n_Not_=59, n_Embedded_=26). **(D)** Difference between subjective scores for easiness to extract information from peer-reviewed to preprint version by embedding of figures in preprints from the paired sample. Difference, 95% C.I.: 0.2 [-0.2, 0.7]. Student’s t test: t=1.00, p=0.32 (n_Not_=38, n_Embedded_=18).

**Supplemental Tables**

**Table S1** – Questionnaire for evaluating quality of reporting. The first 5 sections are applicable to all article categories and compose the general score. The last 4 sections are applicable only to the corresponding category, classified according to the biological model, and compose category-specific scores. Two questions (marked with an asterisk) were inadvertently pre-registered with partial answers representing a score of 1, which was corrected to 0.5 during analysis of the independent samples comparison. Evaluations of the paired sample took place after this change.

**Title/Abstract**

1. Is the biological model / species of animal under study reported?

(Yes – 1; No – 0)

**Risk of Bias**

1. Do the authors report their funding source(s)?

(Yes – 1; No – 0)

2. Is there a statement describing the presence or absence of conflict of interest?

(Yes, and the statement reports a conflict of interest – 1; Yes, and the statement reports no conflict of interest – 1; No statement is present – 0)

3. Is a sample size calculation reported?

(Sample size calculation is reported, with parameters – 1; Sample size calculation is reported, without parameters – 0.5; Sample size calculation is NOT reported – 0)

4. Is assessment of outcome measures reported to be done in a blinded fashion?

(Yes (blinded) – 1; No – 0; Automated/Not applicable)

**Drugs and Reagents**

1. Are the suppliers for all drugs or other treatments in the data under analysis reported?

(Yes – 1; Partially – 0.5; No – 0; Not applicable)

2. Is every antibody used in the data under analysis linked to a citation, catalogue number, clone number or validation profile?

(Yes – 1; No – 0; Not applicable)

3. For pharmacological interventions, is the dose/concentration reported?

(Yes – 1; Partially – 0.5; No – 0; Not applicable)

4. For pharmacological interventions, is the vehicle reported?

(Yes – 1; No – 0; Not applicable)

**Data Presentation**

1. Are the groups compared clearly described?

(Yes – 1; No – 0)

2. Does the study provide a clear timeline for the experimental procedures or exposures and the measurement of outcomes in the data under analysis?

(Yes – 1; No – 0; Not applicable)

3. Is a well-defined summary estimate (e.g. mean or median) of quantitative variables provided for each group? (If “Not applicable” is chosen, please provide the reason)

(Yes – 1; No – 0; Not applicable)

4. Are findings presented with a well-defined measure of variation or precision (e.g. SD/SEM/X%CI)? (If “Not applicable” is chosen, please provide the reason)

(Yes – 1; No – 0; Not applicable)

5. Are unit level data presented?

(Yes (in figures) – 1; Yes (in raw data) – 1 ; Yes (both) – 1; No – 0; Not applicable)

6. Are all data shown in figures or tables clearly attributable to a specific experimental group/condition?

(Yes – 1; No – 0; Not applicable)

7. Are the units for each quantitative measure/indexes shown in figures clearly described?

(Yes – 1; No – 0; Not applicable)

8. Is the meaning of any symbols used in figures/tables (e.g. *, #, ^a^) clearly described?

(Yes – 1; No – 0; Not applicable)

**Data Analysis**

1. Is the experimental unit used for analysis clear?

(Yes – 1; No – 0; Not applicable)

2. Is sample size reported for each group?

(Yes (exact) – 1; Yes (range) – 0.5*; Partially – 0.5; No – 0)

3. Are the statistical tests used clearly described?

(Yes – 1; No – 0)

4. Are the variables and groups to which each statistical result refers to made clear?

(Yes – 1; No – 0)

5. Are the results of statistical tests in the figure (including omnibus and post-hoc comparisons) provided (as a p value or otherwise)?

(Yes – 1; Partially – 0.5; No – 0)

6. Are exact p values reported up to 2 decimal units (e.g. p=0.46, p=0.05, p<0.01)?

(Yes – 1; Partially – 0.5; No – 0)

***In vitro* studies**

1. Was the source of cell lines or microorganisms provided?

(Yes – 1; Partially – 0.5; No – 0; Not applicable)

2. For studies involving cell lines or microorganisms, do the authors report whether they have been authenticated recently (e.g., by STR profiling, within 1 year of use)?

(Yes – 1; Yes, but timing is not mentioned (or is more than one year before experiments) – 0.5; Partially – 0.5; No – 0; Not applicable)

3. Is the culture medium reported?

(Yes – 1; Partially – 0.5; No – 0)

4. Are the culture conditions (temperature, [CO_2_] and presence of O_2_) reported?

(Yes – 1; Partially – 0.5; No – 0)

**Animal studies (invertebrates)**

1. Is the animal species reported? (if yes, please specify which species)

(Yes – 1; No – 0)

2. Is the strain of the animals reported?

(Yes – 1; No – 0; Not applicable)

3. Is the sex of the animals reported?

(Yes – 1; No – 0; Not applicable)

4. Is the age of the animals reported?

(Yes (exact) – 1; Yes (range) – 0.5; No – 0; Not applicable)

5. Is the source/supplier of the animals reported?

(Yes – 1; No – 0; Not applicable)

6. For *in vivo* pharmacological interventions, is the route of administration reported?

(Yes – 1; No – 0; Not applicable)

7. If anaesthesia was performed, are type, route and dose/concentration described?

(Yes – 1; No – 0; Not applicable)

8. Is the method of euthanasia/tissue collection reported?

(Yes – 1; No – 0; Not applicable)

**Animal studies (vertebrates)**

1. Is the animal species reported? (If yes, please specify which species)

(Yes – 1; No – 0)

2. Is the strain of the animals reported?

(Yes – 1; No – 0; Not applicable)

3. Is the sex of the animals reported?

(Yes – 1; No – 0; Not applicable)

4. Is the age of the animals reported?

(Yes (exact) – 1; Yes (range) – 0.5; No – 0; Not applicable)

5. Is the number of animals housed together reported?

(Yes (exact) – 1; Yes (range) – 0.5; No – 0; Not applicable)

6. Is the source/supplier of the animals reported?

(Yes – 1; No – 0; Not applicable)

7. Are animals reported to be randomized to experimental groups?

(Yes – 1; No – 0; Not applicable)

8. For *in vivo* pharmacological interventions, is the route of administration reported?

(Yes – 1; No – 0; Not applicable)

9. If anaesthesia was performed, are type, route and dose/concentration described?

(Yes – 1; No – 0; Not applicable)

10. Is the method of euthanasia/tissue collection reported?

(Yes – 1; No – 0; Not applicable)

11. Does the manuscript include an explicit statement of approval by a clearly identified ethics committee?

(Yes (includes approval and committee) – 1; Yes (includes approval but no committee) – 0.5*; No – 0; Not applicable)

12. Does the manuscript name the international, national or institutional guidelines followed?

(Yes – 1; No – 0; Not applicable)

**Human studies**

1. Does the manuscript describe the recruitment process (including the target population)?

(Yes – 1; No – 0)

2. Are the eligibility criteria adequately described?

(Yes – 1; No – 0)

3. Is the sex of the subjects reported?

(Yes – 1; No – 0; Not applicable)

4. Is the age range of the subjects reported?

(Yes – 1; No – 0)

5. Are subjects reported to be randomized to experimental groups?

(Yes – 1; No – 0; Not applicable)

6. Are the subjects reported to be blinded to the experimental group?

(Yes – 1; No – 0; Not applicable)

7. For pharmacological interventions, is the route of administration reported?

(Yes – 1; No – 0; Not applicable)

8. Does the manuscript include an explicit statement of ethical approval and identify the committee(s) approving the study protocol?

(Yes (includes approval and committee)– 1; Yes (includes approval but no committee) – 0.5; No – 0)

9. Does the manuscript name the international, national or institutional guidelines followed?

(Yes – 1; No – 0)

10. Does the manuscript report that every subject signed an informed consent form?

(Yes – 1; No – 0; Not applicable)

**Table S2** – Inter-evaluator agreement on all analyses. The top part of the table shows mean agreement between each pair of evaluators (labelled as A to Q) in both independent and paired samples of articles. The bottom part of the table shows the number of articles analyzed by each pair of evaluators. The last column shows the mean agreement of each evaluator with all others.

|  | **A** | **B** | **C** | **D** | **E** | **F** | **G** | **H** | **I** | **J** | **K** | **L** | **M** | **N** | **O** | **P** | **Q** | **Mean** |
| --- | --- | --- | --- | --- | --- | --- | --- | --- | --- | --- | --- | --- | --- | --- | --- | --- | --- | --- |
| **A** | - | 76.9% | 77.6% | 79.1% | 78.5% | 77.2% | 74.5% | - | 73.0% | 74.8% | 76.8% | 74.8% | 82.1% | 78.2% | 78.8 % | - | - | 77.1% |
| **B** | 12 | - | 79.8% | 84.4% | 81.8% | 72.2% | 79.3% | 78.5% | 83.9% | 78.3% | 81.8% | 78.4% | 81.1% | - | 82.1% | 81.8% | 80.5% | 80.1% |
| **C** | 9 | 7 | - | 77.0% | 80.6% | 75.6% | 76.4% | 81.4% | 66.7% | 80.2% | 76.5% | 70.8% | 81.6% | 79.6% | 84.8% | 78.9% | 86.3% | 78.4% |
| **D** | 9 | 20 | 7 | - | 81.9% | 67.7% | 75.4% | 78.9% | 75.7% | 85.1% | 81.4% | 82.0% | 81.6% | 81.8% | 86.8% | 85.2% | - | 80.3% |
| **E** | 10 | 10 | 9 | 13 | - | 76.1% | 79.0% | 84.3% | 82.9% | 84.5% | 81.8% | 76.5% | 78.7% | 82.7% | 80.8% | - | 88.9% | 81.3% |
| **F** | 7 | 8 | 7 | 4 | 4 | - | 76.0% | 74.5% | 71.4% | 74.1% | 69.1% | 71.8% | 74.3% | 72.6% | 70.1% | 73.0% | 84.8% | 73.8% |
| **G** | 4 | 7 | 7 | 9 | 8 | 6 | - | 75.2% | 71.3% | 77.4% | 80.4% | 75.0% | 82.3% | 77.4% | 79.4% | 77.2% | 70.4% | 76.7% |
| **H** | 0 | 4 | 9 | 8 | 6 | 3 | 5 | - | 78.6% | 82.8% | 88.3% | 82.7% | 83.0% | - | 79.3% | - | - | 80.6% |
| **I** | 2 | 2 | 3 | 7 | 5 | 6 | 7 | 6 | - | 78.6% | 87.8% | 80.2% | 82.9% | 83.2% | 85.4% | - | - | 78.7% |
| **J** | 12 | 4 | 12 | 3 | 4 | 11 | 5 | 5 | 6 | - | 75.9% | 81.3% | 79.4% | 83.8% | 81.2% | 74.2% | 86.6% | 79.9% |
| **K** | 8 | 9 | 4 | 5 | 5 | 3 | 4 | 4 | 2 | 7 | - | 82.0% | 83.3% | 80.0% | 80.8% | - | 81.1% | 80.5% |
| **L** | 7 | 2 | 9 | 8 | 3 | 16 | 8 | 2 | 3 | 11 | 9 | - | 81.1% | 86.9% | 83.9% | 81.2% | - | 79.2% |
| **M** | 4 | 12 | 6 | 9 | 6 | 2 | 8 | 4 | 2 | 9 | 7 | 9 | - | 84.7% | 80.7% | 81.3% | - | 81.2% |
| **N** | 7 | 4 | 6 | 1 | 6 | 11 | 9 | 0 | 8 | 9 | 5 | 9 | 8 | - | 86.2% | 86.4% | 81.5% | 81.8% |
| **O** | 3 | 16 | 5 | 10 | 4 | 9 | 8 | 4 | 2 | 5 | 4 | 14 | 14 | 9 | - | 87.9% | 77.8% | 81.6% |
| **P** | 0 | 1 | 4 | 1 | 0 | 6 | 3 | 0 | 0 | 4 | 0 | 1 | 2 | 6 | 1 |  | 84.6% | 81.1% |
| **Q** | 0 | 2 | 4 | 0 | 2 | 1 | 2 | 0 | 0 | 3 | 2 | 0 | 0 | 4 | 1 | 3 |  | 82.3% |

**Table S3** – Individual evaluators’ mean overall scores per group in both stages of the project. Results are presented as Mean ± S.D. (number of articles). Two-way ANOVA shows that there was no interaction between evaluator and group in either sample (F=1.28, p_Interaction_=0.22 for the independent sample; F=1.05, p_Interaction_=0.40 for the paired sample).

|  | **Independent sample** | | **Paired sample** | |
| --- | --- | --- | --- | --- |
| **Evaluator** | **bioRxiv** | **PubMed** | **Preprint** | **Peer-Reviewed** |
| A | 59.3±12.4 (16) | 64.8±11.4 (16) | 56.2±8.3 (6) | 63.2±10.5 (9) |
| B | 68.3±9.6 (16) | 68.0±10.9 (16) | 67.5±11.7 (17) | 69.1±12.7 (12) |
| C | 66.4±17.5 (15) | 66.6±11.8 (13) | 64.3±13.4 (12) | 72.0±9.3 (13) |
| D | 65.5±13.4 (16) | 72.4±11.6 (16) | 69.9±10.3 (12) | 71.4±8.2 (14) |
| E | 68.5±11.9 (14) | 66.6±11.1 (14) | 63.5±10.6 (11) | 74.4±7.1 (8) |
| F | 62.5±8.4 (14) | 67.8±9.2 (15) | 62.9±10.8 (12) | 58.6±9.3 (10) |
| G | 66.7±10.1 (13) | 72.0±11.8 (14) | 67.7±8.8 (13) | 69.4±7.7 (13) |
| H | 64.6±10.1 (15) | 66.4±10.8 (16) | - | - |
| I | 65.9±15.3 (16) | 65.5±15.2 (14) | 65.6 (1) | 82.6 (1) |
| J | 65.6±10.0 (14) | 67.1±7.6 (13) | 63.4±10.5 (13) | 72.0±10.9 (15) |
| K | 64.6±14.5 (14) | 74.7±7.2 (15) | 66.6±4.1 (4) | 63.6±12.7 (7) |
| L | 63.6±10.5 (14) | 75.6±8.3 (14) | 64.0±12.6 (13) | 74.3±10.6 (15) |
| M | 66.1±8.0 (11) | 77.2±8.6 (14) | 65.6±12.9 (13) | 71.8±11.0 (13) |
| N | 68.0±11.0 (14) | 68.0±9.8 (12) | 65.8±10.6 (12) | 72.1±6.3 (13) |
| O | 65.5±10.1 (14) | 76.2±9.0 (14) | 68.8±9.9 (14) | 75.7±13.0 (12) |
| P | - | - | 69.5±12.5 (8) | 68.8±1.7 (8) |
| Q | - | - | 71.2±10.6 (7) | 70.4±8.9 (5) |

**Table S4** - Complete sample description. Columns refer to bioRxiv and PubMed articles included in the independent samples comparison and to preprint/published article pairs included in the paired comparison (in which preprints partially overlap with the bioRxiv independent sample). Subject areas for preprints were extracted from bioRxiv, whereas PubMed articles were categorized by two evaluators in one of the bioRxiv categories. In the paired sample, the area was extracted only from bioRxiv, while region of origin and animal species did not differ between preprint and peer-reviewed versions. One peer-reviewed article and one preprint did not report animal species while their counterpart did. For simplicity, we show the reported species for the pair.

| **Region of origin** | **bioRxiv** | **PubMed** | **Pairs** |
| --- | --- | --- | --- |
| Africa | 0 0 % | 2 2.6 % | 0 0 % |
| Asia | 5 6.6 % | 18 23.7 % | 2 3.6 % |
| Europe | 32 42.1 % | 27 35.5 % | 21 37.5 % |
| Latin America | 1 1.3 % | 4 5.3 % | 0 0 % |
| North America | 34 44.7 % | 23 30.3 % | 31 55.3 % |
| Oceania | 4 5.3 % | 2 2.6 % | 2 3.6 % |
| **Subject Area** |  |  |  |
| Animal Behavior and Cognition | 1 1.3 % | 1 1.3 % | 1 1.8 % |
| Biochemistry | 1 1.3 % | 1 1.3 % | 2 3.6 % |
| Bioengineering | 1 1.3 % | 2 2.6 % | 1 1.8 % |
| Bioinformatics | 2 2.6 % | 1 1.3 % | 0 0 % |
| Cancer Biology | 0 0 % | 3 3.9 % | 0 0 % |
| Cell Biology | 6 7.9 % | 2 2.6 % | 3 5.3 % |
| Clinical Trials | 0 0 % | 9 11.8 % | 0 0 % |
| Developmental Biology | 2 2.6 % | 3 3.9 % | 2 3.6 % |
| Epidemiology | 0 0 % | 9 11.8 % | 0 0 % |
| Evolutionary Biology | 5 6.6 % | 0 0 % | 3 5.3 % |
| Genetics | 5 6.6 % | 1 1.3 % | 7 12.5 % |
| Genomics | 5 6.6 % | 1 1.3 % | 3 5.3 % |
| Immunology | 1 1.3 % | 3 3.9 % | 1 1.8 % |
| Microbiology | 7 9.2 % | 5 6.6 % | 3 5.3 % |
| Molecular Biology | 2 2.6 % | 5 6.6 % | 2 3.6 % |
| Neuroscience | 34 44.7 % | 7 9.2 % | 23 41.1 % |
| Other | 0 0 % | 2 2.6 % | 1 1.8 % |
| Pathology | 0 0 % | 2 2.6 % | 0 0 % |
| Pharmacology and Toxicology | 0 0 % | 12 15.8 % | 0 0 % |
| Physiology | 1 1.3 % | 6 7.9 % | 1 1.8 % |
| Scientific Communication and Education | 0 0 % | 1 1.3 % | 0 0 % |
| Synthetic Biology | 1 1.3 % | 0 0 % | 1 1.8 % |
| Systems Biology | 2 2.6 % | 0 0 % | 2 3.6 % |
| **Species (invertebrates)** |  |  |  |
| *Daphnia magna* | 0 0 % | 1 100 % | 0 0 % |
| *Drosophila* sp. | 1 100 % | 0 0 % | 5 55.6 % |
| *C. elegans* | 0 0 % | 0 0 % | 3 33.3 % |
| *D. melanogaster* & *A. gambiae* | 0 0 % | 0 0 % | 1 11.1 % |
| **Species (vertebrates)** |  |  |  |
| Atlantic Cod | 0 0 % | 1 4 % | 0 0 % |
| Buffalo | 0 0 % | 1 4 % | 0 0 % |
| Chicken | 2 8 % | 0 0 % | 1 7.1 % |
| Cow | 0 0 % | 1 4 % | 0 0 % |
| *Macaca* sp. | 3 12 % | 0 0 % | 2 14.3 % |
| Mice | 14 56 % | 7 28 % | 7 50 % |
| *Peromyscus eremicus* | 0 0 % | 0 0 % | 1 7.1 % |
| Pig | 0 0 % | 2 8 % | 0 0 % |
| Porpoise | 1 4 % | 0 0 % | 0 0 % |
| Rat | 1 4 % | 12 48 % | 1 7.1 % |
| Turtle | 1 4 % | 0 0 % | 1 7.1 % |
| Zebrafish | 3 12 % | 1 4 % | 1 7.1 % |

**Table S5** – Reporting scores by article category (as defined by biological model). Independent sample comparisons’ results are from Student’s t tests, while paired samples’ results are from paired t tests. Sample size refers to the number of articles per category. 2-way ANOVA results are presented in individual lines below each comparison set. C.I.; Confidence interval of the difference.

| Score | Stage of the study | Subset | Mean ± S.D. (preprints) | Mean ± S.D. (peer-reviewed) | Difference  [95% C.I.] | | p value | Sample Size |
| --- | --- | --- | --- | --- | --- | --- | --- | --- |
| Overall | *Independent sample* | All | 66.9±12.2 | 71.9±10.1 | 5.0 [1.4, 8.6] | | 0.007 | 76 |
|  |  | *In vitro* | 61.2±9.7 | 65.0±7.4 | 3.8 [-2.0, 9.7] | | 0.19 | 18 |
|  |  | Invertebrates | 77.3 | 76.8 | - | | - | 1 |
|  |  | Vertebrates | 66.4±11.1 | 71.6±9.2 | 5.2 [-0.6, 11.0] | | 0.08 | 25 |
|  |  | Humans | 70.1±13.4 | 75.8±10.3 | 5.7 [-0.3, 11.6] | | 0.06 | 32 |
| Group: F=8.30, df=1, p=0.005; Category: F=6.87, df=3, p=0.0002; Interaction: F=0.10, df=3, p=0.96 | | | | | | | | |
| Overall | *Paired sample* | All | 67.6±10.8 | 72.3±10.1 | 4.7 [2.4, 7.0] | | 0.0001 | 56 |
|  |  | *In vitro* | 60.8±8.8 | 66.8±8.7 | 6.1 [1.1, 11.0] | | 0.02 | 14 |
|  |  | Invertebrates | 68.5±7.6 | 73.0±10.1 | 4.5 [-2.4, 11.3] | | 0.17 | 9 |
|  |  | Vertebrates | 66.2±10.0 | 73.5±10.1 | 7.3 [2.2, 12.3] | | 0.008 | 14 |
|  |  | Humans | 73.3±11.3 | 75.2±10.3 | 2.0 [-1.9, 5.8] | | 0.30 | 19 |
| Group: F=6.41, df=1, p=0.013; Category: F=6.06, df=3, p=0.0008; Interaction: F=0.45, df=3, p=0.72 | | | | | | | | |
| General | *Independent sample* | All | 71.0±12.0 | 73.4±10.0 | 2.4 [-1.1, 6.0] | | 0.17 | 76 |
|  |  | *In vitro* | 64.9±11.6 | 66.7±8.6 | 1.8 [-5.2, 8.7] | | 0.61 | 18 |
|  |  | Invertebrates | 77.8 | 81.0 | - | | - | 1 |
|  |  | Vertebrates | 73.7±11.2 | 74.1±9.8 | 0.4 [-5.5, 6.4] | | 0.88 | 25 |
|  |  | Humans | 72.1±12.2 | 76.5±9.5 | 4.4 [-1.0, 9.9] | | 0.11 | 32 |
| Group: F=2.02, df=1, p=0.16; Category: F=5.89, df=3, p=0.0008; Interaction: F=0.35, df=3, p=0.79 | | | | | | | | |
| General | *Paired sample* | All | 72.6±9.8 | 77.9±9.3 | 5.3 [2.8, 7.7] | | 7.5x10^-5^ | 56 |
|  |  | *In vitro* | 68.1±10.9 | 73.3±9.1 | 5.2 [-0.5, 11.0] | | 0.07 | 14 |
|  |  | Invertebrates | 70.6±6.4 | 76.0±10.5 | 5.3 [-2.1, 12.8] | | 0.13 | 9 |
|  |  | Vertebrates | 75.1±11.3 | 81.7±8.2 | 6.6 [0.2, 12.9] | | 0.04 | 14 |
|  |  | Humans | 75.1±8.1 | 79.4±8.6 | 4.3 [0.6, 7.9] | | 0.02 | 19 |
| Group: F=9.14, df=1, p=0.003; Category: F=4.23, df=3, p=0.007; Interaction: F=0.08, df=3, p=0.97 | | | | | | | | |
| Specific | *Independent sample* | *In vitro* | 44.0±25.0 | 56.2±20.2 | 12.2 [-3.1, 27.7] | | 0.11 | 18 |
|  |  | Invertebrates | 75.0 | 64.3 | - | | - | 1 |
|  |  | Vertebrates | 51.9±18.2 | 66.3±15.7 | 14.4 [4.7, 24.1] | | 0.004 | 25 |
|  |  | Humans | 65.2±26.9 | 74.5±16.7 | 9.3 [-1.8, 20.5] | | 0.10 | 32 |
| Group: F=11.49, df=1, p=0.0009; Category: F=7.35, df=3, p=0.0001; Interaction: F=0.33, df=3, p=0.80 | | | | | | | | |
| Specific | *Paired sample* | *In vitro* | 25.6±22.0 | 35.4±22.4 | 9.8 [-3.2, 22.8] | | 0.13 | 14 |
|  |  | Invertebrates | 60.4±18.0 | 61.1±22.5 | 0.7 [-12.3, 13.7] | | 0.90 | 9 |
|  |  | Vertebrates | 48.2±13.9 | 57.8±21.4 | 9.6 [3.1, 16.2] | | 0.007 | 14 |
|  |  | Humans | 69.2±23.4 | 64.4±23.0 | -4.8[-14.3, 4.7] | 0.30 | | 19 |
| Group: F=0.70, df=1, p=0.41; Category: F=16.44, df=3, p=8.15x10^-9^; Interaction: F=0.90, df=3, p=0.44 | | | | | | | | |

**Table S6** – Reporting of individual questions in the independent samples comparison. Results are for Fisher’s exact test for count data. Only the answers for questions rated as applicable were included in the analysis, causing sample size to vary from question to question. C.I.; confidence interval.

|  | |  | **Yes** | **Part.** | **No** | **Odds ratio [95%C.I.]** | **p value** |
| --- | --- | --- | --- | --- | --- | --- | --- |
|  | **Title/Abstract** | | | | | | |
| Biological model/species | | *bioRxiv* | 64  84.2% | - | 12  15.8% | 2.6 [0.8, 10.1] | 0.12 |
|  |  | *PubMed* | 71  93.4% | - | 5  6.6% |  |  |
|  | **Risk of Bias** | | | | | | |
| Funding Source | | *bioRxiv* | 63  82.9% | - | 13  17.1% | 0.7 [0.3, 1.6] | 0.42 |
|  |  | *PubMed* | 58  76.3% | - | 18  23.7% |  |  |
| Conflict of interest statement | | *bioRxiv* | 34  44.7% | - | 42  55.3% | 2.4 [1.2, 4.8] | 0.01 |
|  |  | *PubMed* | 50  65.8% | - | 26  34.2% |  |  |
| Sample size calculation | | *bioRxiv* | 7  9.2% | 1  1.3% | 68  89.5% | - | 0.10 |
|  |  | *PubMed* | 2  2.6% | 0  0% | 74  97.4% |  |  |
| Blinded assessment of outcomes | | *bioRxiv* | 2  4.1% | - | 47  95.9% | 1.3 [0.1, 15.9] | 1.00 |
|  |  | *PubMed* | 3  5.2% | - | 55  94.8% |  |  |
|  | **Drugs and Reagents** | | | | | | |
| Suppliers | | *bioRxiv* | 13  48.1% | 5  18.5% | 9  33.4% | - | 0.02 |
|  |  | *PubMed* | 31  81.6% | 3  7.9% | 4  10.5% |  |  |
| Antibody validation | | *bioRxiv* | 7  53.8% | - | 6  46.2% | ∞ [0.2, ∞] | 0.25 |
|  |  | *PubMed* | 3  100% | - | 0  0% |  |  |
| Dose/concentration (for pharmacological interventions) | | *bioRxiv* | 18  94.7% | 0  0% | 1  5.3% | - | 1.00 |
|  |  | *PubMed* | 27  87.1% | 1  3.2% | 3  9.7% |  |  |
| Vehicle (for pharmacological interventions) | | *bioRxiv* | 11  68.7% | - | 5  31.3% | 0.75 [0.2, 3.2] | 0.75 |
|  |  | *PubMed* | 18  62.1% | - | 11  37.9% |  |  |
|  | **Data presentation** | | | | | | |
| Clear description of groups | | *bioRxiv* | 74  97.4% | - | 2  2.6% | 2.0 [0.1, 121.1] | 1.00 |
|  |  | *PubMed* | 75  98.7% | - | 1  1.3% |  |  |
| Clear timeline | | *bioRxiv* | 59  84.3% | - | 11  15.7% | 1.1 [0.4, 3.1] | 1.00 |
|  |  | *PubMed* | 59  85.5% | - | 10  14.5% |  |  |
| Summary estimate definition | | *bioRxiv* | 48  65.7% | - | 25  34.3% | 2.3 [1.0, 5.4] | 0.04 |
|  |  | *PubMed* | 57  81.4% | - | 13  18.6% |  |  |
| Variation/precision measure definition | | *bioRxiv* | 49  66.2% | - | 25  33.8% | 2.8 [1.2, 6.9] | 0.01 |
|  |  | *PubMed* | 60  84.5% | - | 11  15.5% |  |  |
| Unit level data | | *bioRxiv* | 22  29% | - | 54  71% | 0.1 [0.02, 0.4] | 4.5x10^-5^ |
|  |  | *PubMed* | 3  4.2% | - | 69  95.8% |  |  |
| Clear group attribution of data | | *bioRxiv* | 76  100% | - | 0  0% | - | - |
|  |  | *PubMed* | 76  100% | - | 0  0% |  |  |
| Units description | | *bioRxiv* | 65  86.7% | - | 10  13.3% | 2.0 [0.6, 8.1] | 0.27 |
|  |  | *PubMed* | 67  93.1% | - | 5  6.9% |  |  |
| Symbol meaning | | *bioRxiv* | 27  69.2% | - | 12  30.8% | 4.9 [1.3, 23.0] | 0.01 |
|  |  | *PubMed* | 45  91.8% | - | 4  8.2% |  |  |
|  | **Data analysis** | | | | | | |
| Experimental unit | | *bioRxiv* | 72  94.7% | - | 4  5.3% | 0.3 [0.1, 1.2] | 0.10 |
|  |  | *PubMed* | 65  85.5% | - | 11  14.5% |  |  |
| Exact sample size description | | *bioRxiv* | 62  81.6% | 5  6.6% | 9  11.8% | - | 0.57 |
|  |  | *PubMed* | 58  76.4% | 9  11.8% | 9  11.8% |  |  |
| Statistical tests used | | *bioRxiv* | 62  81.6% | - | 14  18.4% | 1.7 [0.6, 4.7] | 0.37 |
|  |  | *PubMed* | 67  88.2% | - | 9  11.8% |  |  |
| Variables and groups compared | | *bioRxiv* | 69  90.8% | - | 7  9.2% | 1.8 [0.4, 8.9] | 0.53 |
|  |  | *PubMed* | 72  94.7% | - | 4  5.3% |  |  |
| Complete statistical results | | *bioRxiv* | 58  76.3% | 7  9.2% | 11  15.5% | - | 0.06 |
|  |  | *PubMed* | 49  64.5% | 18  23.7% | 9  11.8% |  |  |
| Exact p values | | *bioRxiv* | 49  64.5% | 6  7.9% | 21  27.6% | - | 0.18 |
|  |  | *PubMed* | 39  51.3% | 12  15.8% | 25  32.9% |  |  |
|  | **In vitro studies** | | | | | | |
| Source/Supplier | | *bioRxiv* | 9  50% | 2  11.1% | 7  38.9% | - | 0.43 |
|  |  | *PubMed* | 12  66.7% | 0  0% | 6  33.3% |  |  |
| Cell line authentication | | *bioRxiv* | 0  0% | 0  0% | 16  100% | - | - |
|  |  | *PubMed* | 0  0% | 0  0% | 18  100% |  |  |
| Culture medium | | *bioRxiv* | 14  77.8% | 0  0% | 4  22.2% | 4.7 [0.4, 252.6] | 0.34 |
|  |  | *PubMed* | 17  94.4% | 0  0% | 1  5.6% |  |  |
| Culture conditions | | *bioRxiv* | 3  16.7% | 6  33.3% | 9  50% | - | 0.04 |
|  |  | *PubMed* | 7  38.9% | 9  50% | 2  11.1% |  |  |
|  | **Animal studies (invertebrates)** | | | | | | |
| Species | | *bioRxiv* | 1  100% | - | 0  0% | - | - |
|  |  | *PubMed* | 1  100% | - | 0  0% |  |  |
| Strain | | *bioRxiv* | - | - | - | - | - |
|  |  | *PubMed* | 0  0% | - | 1  100% |  |  |
| Sex | | *bioRxiv* | 1  100% | - | 0  0% | 0 [0, 39] | 1.00 |
|  |  | *PubMed* | 0  0% | - | 1  100% |  |  |
| Age | | *bioRxiv* | - | - | - | - | - |
|  |  | *PubMed* | 0  0% | 1  100% | 0  0% |  |  |
| Source/supplier | | *bioRxiv* | 1  100% | - | 0  0% | - | - |
|  |  | *PubMed* | 1  100% | - | 0  0% |  |  |
| Route of administration (for pharmacological interventions) | | *bioRxiv* | - | - | - | - | - |
|  |  | *PubMed* | 1  100% | - | 0  0% |  |  |
| Anesthesia description | | *bioRxiv* | - | - | - | - | - |
|  |  | *PubMed* | - | - | - |  |  |
| Euthanasia/Tissue collection method | | *bioRxiv* | 0  0% | - | 1  100% | ∞ [0.03, ∞] | 1.00 |
|  |  | *PubMed* | 1  100% | - | 0  0% |  |  |
|  | **Animal studies (vertebrates)** | | | | | | |
| Species | | *bioRxiv* | 25  100% | - | 0  0% | - | - |
|  |  | *PubMed* | 25  100% | - | 0  0% |  |  |
| Strain | | *bioRxiv* | 18  72% | - | 7  28% | 9.0 [1.0, 436.7] | 0.05 |
|  |  | *PubMed* | 24  96% | - | 1  4% |  |  |
| Sex | | *bioRxiv* | 12  50% | - | 12  50% | 2.9 [0.8, 12.4] | 0.13 |
|  |  | *PubMed* | 18  75% | - | 6  25% |  |  |
| Age | | *bioRxiv* | 5  20.8% | 9  37.5% | 10  41.7% | - | 0.72 |
|  |  | *PubMed* | 7  29.2% | 7  29.2% | 10  41.6% |  |  |
| Housing number | | *bioRxiv* | 1  4.3% | 1  4.3% | 21  91.4% | - | 0.60 |
|  |  | *PubMed* | 3  12.5% | 2  8.3% | 19  79.2% |  |  |
| Source/supplier | | *bioRxiv* | 12  48% | - | 13  52% | 7.6 [1.6, 49.9] | 0.005 |
|  |  | *PubMed* | 22  88% | - | 3  22% |  |  |
| Randomization | | *bioRxiv* | 0  0% | - | 14  100% | ∞ [2.0, ∞] | 0.003 |
|  |  | *PubMed* | 8  47% | - | 9  53% |  |  |
| Route of administration (for pharmacological interventions) | | *bioRxiv* | 6  100% | - | 0  0% | - | - |
|  |  | *PubMed* | 10  100% | - | 0  0% |  |  |
| Anesthesia description | | *bioRxiv* | 10  66.7% | - | 5  33.3% | 0.8 [0.1, 5.5] | 1.00 |
|  |  | *PubMed* | 6  60% | - | 4  40% |  |  |
| Euthanasia | | *bioRxiv* | 8  47% | - | 9  53% | 0.9 [0.2, 4.1] | 1.00 |
|  |  | *PubMed* | 8  44.4% | - | 10  55.6% |  |  |
| Approval by ethics committee | | *bioRxiv* | 14  56% | 0  0% | 11  44% | 1.6 [0.4, 5.9] | 0.56 |
|  |  | *PubMed* | 16  66.7% | 0  0% | 8  33.3% |  |  |
| Ethics guidelines followed | | *bioRxiv* | 14  58.3% | - | 10  41.7% | 1.5 [0.4, 5.7] | 0.56 |
|  |  | *PubMed* | 17  68% | - | 8  32% |  |  |
|  | **Human studies** | | | | | | |
| Recruitment process | | *bioRxiv* | 19  59.4% | - | 13  40.6% | 4.7 [1.2, 22.7] | 0.02 |
|  |  | *PubMed* | 28  87.5% | - | 4  12.5% |  |  |
| Eligibility criteria | | *bioRxiv* | 19  59.4% | - | 13  40.6% | 6.4 [1.5, 39.8] | 0.008 |
|  |  | *PubMed* | 29  90.6% | - | 3  9.4% |  |  |
| Sex | | *bioRxiv* | 27  87.1% | - | 4  12.9% | 1.4 [0.2, 10.6] | 0.71 |
|  |  | *PubMed* | 29  90.6% | - | 3  9.4% |  |  |
| Age range | | *bioRxiv* | 26  81.2% | - | 6  18.8% | 1.2 [0.3, 5.8] | 1.00 |
|  |  | *PubMed* | 27  84.4% | - | 5  15.6% |  |  |
| Randomization | | *bioRxiv* | 1  25% | - | 3  75% | 16.1 [0.7, 1388.8] | 0.05 |
|  |  | *PubMed* | 8  88.9% | - | 1  11.1% |  |  |
| Blinding of subjects | | *bioRxiv* | 1  14.3% | - | 6  85.7% | 1.2 [0.01, 109.6] | 1.00 |
|  |  | *PubMed* | 1  16.7% | - | 5  83.3% |  |  |
| Route of administration (for pharmacological interventions) | | *bioRxiv* | 2  100% | - | 0  0% | 0 [0, 58.4] | 1.00 |
|  |  | *PubMed* | 2  66.7% | - | 1  33.3% |  |  |
| Ethics committee approval | | *bioRxiv* | 23  71.9% | 0  0% | 9  28.1% | - | 0.45 |
|  |  | *PubMed* | 23  71.9% | 2  6.2% | 7  21.9% |  |  |
| Ethics guidelines followed | | *bioRxiv* | 8  25% | - | 24  75% | 1.3 [0.4, 4.7] | 0.78 |
|  |  | *PubMed* | 10  31.2% | - | 22  68.8% |  |  |
| Signed informed consent | | *bioRxiv* | 27  90% | - | 3  10% | 0.2 [0.03, 1.3] | 0.07 |
|  |  | *PubMed* | 15  68.2% | - | 7  31.8% |  |  |

**Table S7** – Reporting of individual questions in the paired samples. Results are for McNemar’s exact test. Only the answers for questions rated as applicable are shown; thus, sample sizes vary for each individual question.

|  | |  | **Yes** | **Part.** | **No** | **Odds ratio [95% C.I.]** | **p value** |
| --- | --- | --- | --- | --- | --- | --- | --- |
|  | **Title/Abstract** | | | | | | |
| Biological model/species | | *bioRxiv* | 48  85.7% | - | 8  14.3% | 0.5 [0.008, 9.6] | 1.00 |
|  |  | *Peer-reviewed* | 47  83.9% | - | 9  16.1% |  |  |
|  | **Risk of Bias** | | | | | | |
| Funding Source | | *bioRxiv* | 50  89.3% | - | 6  10.7% | ∞ [0.9, ∞] | 0.06 |
|  |  | *Peer-reviewed* | 55  98.2% | - | 1  1.8% |  |  |
| Conflict of interest statement | | *bioRxiv* | 26  46.4% | - | 30  53.6% | 11.5 [2.8, 100.6] | 1.9x10^-5^ |
|  |  | *Peer-reviewed* | 47  83.9% | - | 9  16.1% |  |  |
| Sample size calculation | | *bioRxiv* | 4  7.1% | 1  1.8% | 51  91.1% | - | - |
|  |  | *Peer-reviewed* | 6  10.7% | 1  1.8% | 49  87.5% |  |  |
| Blinded assessment of outcomes | | *bioRxiv* | 0  0% | - | 37  100% | - | - |
|  |  | *Peer-reviewed* | 0  0% | - | 37  100% |  |  |
|  | **Drugs and Reagents** | | | | | | |
| Suppliers | | *bioRxiv* | 14  70% | 1  5% | 5  25% | - | - |
|  |  | *Peer-reviewed* | 13  65% | 4  20% | 3  15% |  |  |
| Antibody validation | | *bioRxiv* | 3  42.9% | - | 4  57.1% | - | 1.00 |
|  |  | *Peer-reviewed* | 4  66.7% | - | 2  33.3% |  |  |
| Dose/concentration (for pharmacological interventions) | | *bioRxiv* | 13  81.25% | 1  6.25% | 2  12.5% | - | - |
|  |  | *Peer-reviewed* | 9  75% | 0  0% | 3  25% |  |  |
| Vehicle (for pharmacological interventions) | | *bioRxiv* | 5  41.7% | - | 7  58.3% | ∞ [0.2, ∞] | 0.50 |
|  |  | *Peer-reviewed* | 8  72.7% | - | 3  27.3% |  |  |
|  | **Data presentation** | | | | | | |
| Clear description of groups | | *bioRxiv* | 56  100% | - | 0  0% | - | - |
|  |  | *Peer-reviewed* | 56  100% | - | 0  0% |  |  |
| Clear timeline | | *bioRxiv* | 47  87% | - | 7  13% | 2.5 [0.4, 26.5] | 0.45 |
|  |  | *Peer-reviewed* | 50  92.6% | - | 4  7.4% |  |  |
| Summary estimate definition | | *bioRxiv* | 40  75.5% | - | 13  24.5% | 0.9 [0.2, 3.0] | 1.00 |
|  |  | *Peer-reviewed* | 38  71.7% | - | 15  28.3% |  |  |
| Variation/precision measure definition | | *bioRxiv* | 34  64.1% | - | 19  35.9% | 6.0 [0.7, 276.0] | 0.12 |
|  |  | *Peer-reviewed* | 38  71.7% | - | 15  28.3% |  |  |
| Unit level data | | *bioRxiv* | 14  25% | - | 42  75% | ∞ [0.9, ∞] | 0.06 |
|  |  | *Peer-reviewed* | 19  33.9% | - | 37  66.1% |  |  |
| Clear group attribution of data | | *bioRxiv* | 56  100% | - | 0  0% | - | - |
|  |  | *Peer-reviewed* | 56  100% | - | 0  0% |  |  |
| Units description | | *bioRxiv* | 53  96.4% | - | 2  3.6% | 0.7 [0.05, 5.8] | 1.00 |
|  |  | *Peer-reviewed* | 52  94.5% | - | 3  5.5% |  |  |
| Symbol meaning | | *bioRxiv* | 21  84% | - | 4  16% | ∞ [0.2, ∞] | 0.50 |
|  |  | *Peer-reviewed* | 22  91.7% | - | 2  8.3% |  |  |
|  | **Data analysis** | | | | | | |
| Experimental unit | | *bioRxiv* | 52  92.9% | - | 4  7.1% | 4.0 [0.4, 197.0] | 0.37 |
|  |  | *Peer-reviewed* | 55  98.2% | - | 1  1.8% |  |  |
| Exact sample size description | | *bioRxiv* | 42  75% | 8  14.3% | 6  10.7% | - | - |
|  |  | *Peer-reviewed* | 50  89.4% | 3  5.3% | 3  5.3% |  |  |
| Statistical tests used | | *bioRxiv* | 45  80.4% | - | 11  19.6% | 5.0 [0.6, 236.5] | 0.22 |
|  |  | *Peer-reviewed* | 49  87.5% | - | 7  12.5% |  |  |
| Variables and groups compared | | *bioRxiv* | 52  92.9% | - | 4  7.1% | 1.0 [0.1, 13.8] | 1.00 |
|  |  | *Peer-reviewed* | 52  92.9% | - | 4  7.1% |  |  |
| Complete statistical results | | *bioRxiv* | 46  82.1% | 7  12.5% | 3  5.4% | - | - |
|  |  | *Peer-reviewed* | 47  83.9% | 7  12.5% | 2  3.6% |  |  |
| Exact p values | | *bioRxiv* | 36  64.3% | 7  12.5% | 13  23.2% | - | - |
|  |  | *Peer-reviewed* | 35  62.5% | 6  10.7% | 15  26.8% |  |  |
|  | **In vitro studies** | | | | | | |
| Source/Supplier | | *bioRxiv* | 4  28.6% | 1  7.1% | 9  64.3% | - | - |
|  |  | *Peer-reviewed* | 6  42.9% | 2  14.2% | 6  42.9% |  |  |
| Cell line authentication | | *bioRxiv* | 0  0% | 0  0% | 13  100% | - | - |
|  |  | *Peer-reviewed* | 0  0% | 0  0% | 13  100% |  |  |
| Culture medium | | *bioRxiv* | 7  50% | 0  0% | 7  50% | - | - |
|  |  | *Peer-reviewed* | 8  57.1% | 1  7.1% | 5  35.7% |  |  |
| Culture conditions | | *bioRxiv* | 1  7.1% | 3  21.4% | 10  71.5% | - | - |
|  |  | *Peer-reviewed* | 1  7.1% | 6  42.9% | 7  50% |  |  |
|  | **Animal studies (invertebrates)** | | | | | | |
| Species | | *bioRxiv* | 9  100% | - | 0  0% | - | - |
|  |  | *Peer-reviewed* | 9  100% | - | 0  0% |  |  |
| Strain | | *bioRxiv* | 8  88.9% | - | 1  11.1% | 0 [0, 39.0] | 1.00 |
|  |  | *Peer-reviewed* | 7  77.8% | - | 2  22.2% |  |  |
| Sex | | *bioRxiv* | 2  22.2% | - | 7  77.8% | ∞ [0.03, ∞] | 1.00 |
|  |  | *Peer-reviewed* | 3  33.3% | - | 6  66.7% |  |  |
| Age | | *bioRxiv* | 2  25% | 3  37.5% | 3  37.5% | - | - |
|  |  | *Peer-reviewed* | 3  37.5% | 4  50% | 1  12.5% |  |  |
| Source/supplier | | *bioRxiv* | 5  62.5% | - | 3  37.5% | - | 1.00 |
|  |  | *Peer-reviewed* | 5  62.5% | - | 3  37.5% |  |  |
| Route of administration (for pharmacological interventions) | | *bioRxiv* | - | - | - | - | - |
|  |  | *Peer-reviewed* | 0  0% | - | 1  100% |  |  |
| Euthanasia/Tissue collection method | | *bioRxiv* | 3  50% | - | 3  50% | 0 [0, 39.0] | 1.00 |
|  |  | *Peer-reviewed* | 1  25% | - | 3  75% |  |  |
| Anesthesia description | | *bioRxiv* | - | - | - | - | - |
|  |  | *Peer-reviewed* | - | - | - |  |  |
|  | **Animal studies (vertebrates)** | | | | | | |
| Species | | *bioRxiv* | 13  92.9% | - | 1  7.1% | 1.0 [0.01, 78.5] | 1.00 |
|  |  | *Peer-reviewed* | 13  92.9% | - | 1  7.1% |  |  |
| Strain | | *bioRxiv* | 9  64.3% | - | 5  35.7% | 1.0 [0.01, 78.5] | 1.00 |
|  |  | *Peer-reviewed* | 9  64.3% | - | 5  35.7% |  |  |
| Sex | | *bioRxiv* | 6  50% | - | 6  50% | ∞ [0.2, ∞] | 0.50 |
|  |  | *Peer-reviewed* | 8  61.6% | - | 5  38.5% |  |  |
| Age | | *bioRxiv* | 4  28.6% | 4  28.6% | 6  42.8% | - | - |
|  |  | *Peer-reviewed* | 4  28.6% | 6  42.8% | 4  28.6% |  |  |
| Housing number | | *bioRxiv* | 1  7.7% | 0  0% | 12  92.3% | - | 1.00 |
|  |  | *Peer-reviewed* | 1  7.7% | 0  0% | 12  92.3% |  |  |
| Source/supplier | | *bioRxiv* | 7  50% | - | 7  50% | 1.0 [0.01, 78.5] | 1.00 |
|  |  | *Peer-reviewed* | 7  50% | - | 7  50% |  |  |
| Randomization | | *bioRxiv* | 0  0% | - | 5  100% | - | - |
|  |  | *Peer-reviewed* | 0  0% | - | 6  100% |  |  |
| Route of administration (for pharmacological interventions) | | *bioRxiv* | 4  80% | - | 1  20% | - | 1.00 |
|  |  | *Peer-reviewed* | 3  75% | - | 1  25% |  |  |
| Anesthesia description | | *bioRxiv* | 3  33.3% | - | 6  66.7% | - | 1.00 |
|  |  | *Peer-reviewed* | 2  33.3% | - | 4  66.7% |  |  |
| Euthanasia | | *bioRxiv* | 3  27.3% | - | 8  72.7% | ∞ [0.7, ∞] | 0.12 |
|  |  | *Peer-reviewed* | 7  63.6% | - | 4  36.4% |  |  |
| Approval by ethics committee | | *bioRxiv* | 7  50% | 1  7.1% | 6  42.9% | - | - |
|  |  | *Peer-reviewed* | 11  84.6% | 0  0% | 2  15.4% |  |  |
| Ethics guidelines followed | | *bioRxiv* | 8  57.1% | - | 6  42.9% | ∞ [0.2, ∞] | 0.50 |
|  |  | *Peer-reviewed* | 10  71.4% | - | 4  28.6% |  |  |
|  | **Human studies** | | | | | | |
| Recruitment process | | *bioRxiv* | 12  63.2% | - | 7  36.8% | 0.2 [0.004, 1.8] | 0.22 |
|  |  | *Peer-reviewed* | 8  42.1% | - | 11  57.9% |  |  |
| Eligibility criteria | | *bioRxiv* | 12  63.2% | - | 7  36.8% | 0.4 [0.1, 1.6] | 0.23 |
|  |  | *Peer-reviewed* | 7  36.8% | - | 12  63.2% |  |  |
| Sex | | *bioRxiv* | 16  84.2% | - | 3  15.8% | - | 1.00 |
|  |  | *Peer-reviewed* | 16  88.9% | - | 2  11.1% |  |  |
| Age range | | *bioRxiv* | 16  84.2% | - | 3  15.8% | 0 [0, 39.0] | 1.00 |
|  |  | *Peer-reviewed* | 15  78.9% | - | 4  21.1% |  |  |
| Randomization | | *bioRxiv* | 0  0% | - | 2  100% | - | - |
|  |  | *Peer-reviewed* | 0  0% | - | 1  100% |  |  |
| Blinding of subjects | | *bioRxiv* | 0  0% | - | 1  100% | - | - |
|  |  | *Peer-reviewed* | 0  0% | - | 2  100% |  |  |
| Route of administration (for pharmacological interventions) | | *bioRxiv* | 1  100% | - | 0  0% | - | - |
|  |  | *Peer-reviewed* | - | - | - |  |  |
| Ethics committee approval | | *bioRxiv* | 15  78.9% | 0  0% | 4  21.1% | 2.0 [0.1, 118.0] | 1.00 |
|  |  | *Peer-reviewed* | 16  84.2% | 0  0% | 3  15.8% |  |  |
| Ethics guidelines followed | | *bioRxiv* | 6  31.6% | - | 13  68.4% | ∞ [0.2, ∞] | 0.50 |
|  |  | *Peer-reviewed* | 8  42.1% | - | 11  57.9% |  |  |
| Signed informed consent | | *bioRxiv* | 16  84.2% | - | 3  15.8% | ∞ [0.03, ∞] | 1.00 |
|  |  | *Peer-reviewed* | 16  88.9% | - | 2  11.1% |  |  |

**Additional Files**

Supplementary File 1.xls

Supplementary File 2 (updated).RMD

Supplementary Text 1.pdf
